# Complex Mutation Profiles in Mismatch Repair and Ribonucleotide Reductase Mutants Reveal Novel Repair Substrate Specificity of MutS Homolog (MSH) Complexes

**DOI:** 10.1101/2021.06.30.450577

**Authors:** Natalie A. Lamb, Jonathan Bard, Raphael Loll-Krippleber, Grant W. Brown, Jennifer A. Surtees

## Abstract

Determining mutation signatures is standard for understanding the etiology of human tumors and informing cancer treatment. Multiple determinants of DNA replication fidelity prevent mutagenesis that leads to carcinogenesis, including the regulation of free deoxyribonucleoside triphosphate (dNTP) pools by ribonucleotide reductase (RNR) and repair of replication errors by the mismatch repair (MMR) system. We identified genetic interactions between *rnr1* alleles that elevate dNTP levels and MMR. We then utilized a targeted deep-sequencing approach to determine mutational signatures associated with MMR pathway defects. By combining *rnr1* and *msh* mutations to increase dNTP levels and alter the mutational load, we uncovered previously unreported specificities of Msh2-Msh3 and Msh2-Msh6. Msh2-Msh3 is uniquely able to direct repair of G/C single base deletions in GC runs, while Msh2-Msh6 specifically directs repair of substitutions at G/C dinucleotides. We also identified broader sequence contexts that influence variant profiles in different genetic backgrounds. Finally, we observed that the mutation profiles in double mutants were not necessarily an additive relationship of mutation profiles in single mutants. Our results have implications for interpreting mutation signatures from human tumors, particularly when MMR is defective.

## Introduction

Cancer is a genetic disease caused by mutation accumulation; DNA replication is an important source of mutation. Replicative polymerases, Polε and Polδ, minimize errors via a highly selective nucleotide binding domain to prevent deoxynucleotide triphosphate (dNTP) misincorporation and an exonuclease domain to proofread and remove errors (Kunkel 2004; Mcculloch AND KUNKEL 2008; ST Charles *et al*. 2015; Ganai AND JOHANSSON 2016). Together, these functions lead to error rates on the order of 10^-7^ (ST Charles *et al*. 2015; Ganai AND JOHANSSON 2016). Appropriate levels and ratios of the four dNTPs are also essential for maintaining high fidelity polymerase function. This has been demonstrated in yeast using mutations in allosteric sites of *RNR1* that alter dNTP pools in different ways. *RNR1* encodes the large subunit of the enzyme that catalyzes the rate-limiting step in dNTP synthesis, ribonucleotide reductase (RNR). Even a modest 2-fold increase above normal levels, as seen in *rnr1D57N*, increased nucleotide misincorporation by DNA polymerases and elevated mutation rates (Chabes *et al*. 2003; Xu *et al*. 2008). More pronounced and skewed elevations in dNTP pools are generated in *rnr1Y285F* and *rnr1Y285A*, which increase dCTP and dTTP 3-fold and 20-fold, respectively (Kumar *et al*. 2010). These increases in dNTP pool levels further compromise replication fidelity (Kumar *et al*. 2010; Kumar *et al*. 2011b; Buckland *et al*. 2014; Watt *et al*. 2016). Cancer cells have increased proliferative nature and thus elevated dNTP pools may be necessary to support accelerated DNA replication (Mathews 2015; Connor *et al*. 2017), which may, in turn, increase mutagenesis, promote molecular evolution and provide a selective advantage to the tumor.

We previously developed a targeted deep sequencing approach to characterize mutation profiles of these three *rnr1* alleles (Lamb *et al*. 2021). The depth of sequencing allowed a more robust and nuanced analysis of mutation spectra than in previous work (Xu *et al*. 2008; Kumar *et al*. 2011b; Buckland *et al*. 2014; Watt *et al*. 2016). We revealed genotype-specific mutation profiles, including mutation spectra, sequence context and nucleotide motifs, with even modest changes in dNTP pools. In addition to changes in the proportion and/or ratios of SNVs, especially increased CG>TA mutations, all three *rnr1* alleles exhibited a shift in the relative distribution of single base deletions toward G/C deletions, which are typically rare events in wild-type backgrounds (Lamb *et al*. 2021). The frequency of single base G/C deletions was particularly elevated in *rnr1Y285A*. We suggested that the effects of altered dNTP pools on mutation profiles should be incorporated in the analysis of mutation signatures in human cancer (Lamb *et al*. 2021).

The variants we observed in the *rnr1* backgrounds are typically substrates for mismatch repair (MMR), which functions as a spell-check, recognizing and directing the repair of errors in replication (Kunkel AND ERIE 2015), thereby increasing the fidelity of replication by an additional 10-1,000 fold (Mcculloch AND KUNKEL 2008; Ganai AND JOHANSSON 2016). We and others have demonstrated combinatorial effects of *rnr1* and *mmr-* alleles on mutation rates (Xu *et al*. 2008; Kumar *et al*. 2011b; Buckland *et al*. 2014; Watt *et al*. 2016), consistent with the prediction that mutations generated with altered *rnr1* are substrates for MMR. Once recognized by a <u>M</u>ut<u>S h</u>omolog (Msh) complex, the structures generated by these errors are targeted for excision. In most eukaryotes, two heterodimeric complexes bind errors at the replication fork: Msh2-Msh6 and Msh2-Msh3, which recognize a broad spectrum of mismatches and insertion/deletion loops (IDLs) with different specificities. The current model of post-replicative repair posits that Msh2-Msh3 recognizes, binds and directs repair of IDLs up to 17 nucleotides long (Sia *et al*. 1997; Jensen *et al*. 2005), while Msh2-Msh6 targets mismatches and IDLs of 1-2 nucleotides. Msh2-Msh3 also has affinity for some mismatches, especially C-C, A-A and (possibly) G-G (Harrington AND KOLODNER 2007; Srivatsan *et al*. 2014).

MMR is deficient in ∼25% of sporadic cancers caused by increased rate of mutagenesis (Mastrocola AND Heinen 2010). Deficiencies in MMR genes *MSH2, MSH6, PMS2* and *MLH1* cause hereditary Lynch syndrome, which leads to a strong predisposition to cancers of the gastrointestinal tract, endometrial cancer and lymphomas, and are defined by microsatellite instability (Pino *et al*. 2009; Heinen 2010). MMR mutations are also implicated in breast and ovarian cancer (Davies *et al*. 2017; Fusco *et al*. 2018). While Msh2-Msh3 has not been directly linked to Lynch syndrome, mutations in *MSH3* lead to cancer predisposition within and outside the gastrointestinal tract (Edelmann *et al*. 2000; Van OERS *et al*. 2014; Adam *et al*. 2016; Morak *et al*. 2017; Santos *et al*. 2018; Valle *et al*. 2019) as well as chemoresistant tumors (Takahashi *et al*. 2011; Park *et al*. 2013; Nogueira *et al*. 2018). Loss of *MSH3* function leads to elevated alterations of selected tetranucleotide repeats, which is distinct from microsatellite instability and has been associated with a number of different cancers, including up to ∼60% of colorectal cancers (Carethers *et al*. 2015). Therefore, while *MSH3* also plays a role in tumorigenesis, its role is distinct from that of Msh2-Msh6.

The multiplicative and synergistic effects of combining defects in both MMR and *RNR1* on mutation rates (*rnr1D57N msh2Δ, rnr1D57N* msh6Δ (Xu *et al*. 2008) and *rnr1Y285A msh2Δ* (Buckland *et al*. 2014; Watt *et al*. 2016)) indicated a genetic interaction between these pathways. We predicted additional pathways would interact genetically with *rnr1* alleles. We performed synthetic genetic array (SGA) screens to identify pathways that interact genetically with all three *rnr1* alleles. We identified a number of pathways involved in DNA metabolism. Most strikingly, in the *rnr1Y285A* SGA screen, we identified essentially the entire MMR pathway, both Msh2-Msh3- and Msh2-Msh6-medated MMR. Therefore, we focused the characterization of rnr1-MMR genetic interactions at the nucleotide level, using our targeted deep sequencing approach ((Xu *et al*. 2008; Kumar *et al*. 2011b; Buckland *et al*. 2014; Lamb *et al*. 2021)) with an eye to developing a mechanistic understanding of mutation signatures that are observed in tumors. At the same time, the altered mutation profiles generated by *rnr1* alleles (Lamb *et al*. 2021) allowed us to evaluate the role of Msh2-Msh3 and Msh2-Msh6 in directing repair of typically rare replication errors. We characterized single *mshΔ* mutants and evaluated combinatorial effects on the mutation profiles when combined with *rnr1* alleles. We identified novel and specific DNA substrates for Msh2-Msh3- versus Msh2-Msh6-mediated MMR and demonstrated that mutation profiles of *rnr1 mshΔ* double mutants were not necessarily additive of the single mutant profiles, which has implications for the analysis of mutation signatures in human cancers.

## Materials and Methods

### Strains and plasmids

All strains used for sequencing and mutator assays were derived from the W303 *RAD5+* background (Table S1). Strains used for the synthetic lethal screens were derived from S288C (Table S1). Construction of *rnr1D57N*, *rnr1Y285F* and *rnr1Y285A*, with and without *pGAL-RNR1*, was described previously (Lamb *ET AL.* 2021). Mismatch repair genes were deleted by amplifying *msh2Δ::kanMX, msh6Δ::kanMX,* and *msh3Δ::kanMX* chromosomal fragments from deletion collection strains utilizing primers A and D specific to each locus (Table S2). PCR products from these strains were used for transformation to replace the endogenous *MSH* gene with the *kanMX* cassette, which confers resistance to the drug, G418. Transformants were selected on YPD plates containing G418 and deletions were confirmed by PCR. We did not generate *msh3Δ* in the *rn1Y285F/A-pGAL-RNR1* backgrounds.

### Measuring mutation rates at *CAN1*

Mutation rates were measured at the *CAN1* locus as previously described (Xu *et al*. 2008; Lamb *et al*. 2021). Briefly, strains were grown on complete media (YPD) until colonies reach 2 mm in size. Colonies were then suspended in 100 μl of 1x TE (10 mM Tris-HCl, pH 7.5; 1 mM EDTA) and diluted 1:10,000. 20 μl of the undiluted colony suspension was plated on SC-ARG + Canavanine and 100 μl of the 10^-4^ dilution was plated on synthetic complete plates lacking arginine. The plates were incubated at 30°C until colonies reached ∼1 mm in size. Colonies were counted and mutation rates and 95% confidence intervals were calculated though FluCalc fluctuation analysis software (Radchenko *et al*. 2018).

### Synthetic Genetic Array Analysis

The genetic screens were performed using SGA technology (Baryshnikova *et al*. 2010). Briefly, query strains carrying each of the *rnr1* alleles were crossed to an ordered array of all the viable yeast deletion strains and an array of temperature sensitive alleles of yeast essential genes. Diploid cells were transferred to a sporulation-inducing medium, after which the germinated spores were selected for the simultaneous presence of the gene deletion and the *rnr1* allele. Colony size was quantified as a measure of fitness, and SGA scores and p-values were calculated as described in (Baryshnikova *et al*. 2010). SGA scores from deletion mutants and ts mutants were merged by scaling the ts screen scores according to the SGA scores of the deletion mutants that are present in the ts allele array. A z-score was calculated for all the genes in each screen, and a cutoff of z=-2 was applied to identify negative genetic interactions. The SGA data is presented in Tables S4-S10.

### Gene Ontology enrichment analysis

GO term analysis was performed using the GO term finder tool (http://go.princeton.edu/) using a P-value cutoff of 0.01 and applying Bonferroni correction, querying biological process enrichment for each gene set. GO term enrichment results were further processed with REViGO (Supek *et al*. 2011) using the “Medium (0.7)” term similarity filter and simRel score as the semantic similarity measure. Terms with a frequency greater than 15% in the REViGO output were eliminated as too general.

### Spatial Analysis of Functional Enrichment

Network annotations were made with the Python implementation of Spatial Analysis of Functional Enrichment (SAFE) ((Baryshnikova 2016); https://github.com/baryshnikova-lab/safepy). The yeast genetic interaction similarity network and its functional domain annotations were obtained from (Costanzo *et al*. 2016).

### Sample preparation and analysis pipeline

A detailed description of sample preparation and the analytical pipeline used for data analysis can be found in (Lamb *et al*. 2021). Briefly, we pooled ∼2000 colonies Can^R^ colonies for each biological replicate and extracted genomic DNA from the pool. Each genotype was represented by at least 4 biological replicates (see Table S21). The *CAN1* gene was amplified by PCR using KAPA HiFi (Roche) in 6 overlapping fragments that were purified using the Zymo ZR-96 DNA Clean-up Kit. Nextera barcode adaptors were added to the amplicons, followed by attachment of Illumina Nextera XT index primers set A (Illumina). Excess adapters were removed using Ampure XP beads (Beckman Coulter). Pooled samples were diluted to 4 nM, denatured using NaOH and loaded onto an Illumina MiSeq sequencing platform (PE300, V3) with 20% PhiX control in two separate runs to increase coverage and as a check for reproducibility. Paired end (2x300) deep sequencing of *CAN1* provided enough sequencing depth to determine mutation spectra for 150 unique samples, representing over 30 different genotypes, and including biological and technical replicates. *CAN1* was sequenced at an average depth of approximately 16,000 reads per base in *CAN1* per sample allowing for detailed characterization of mutation spectra.

Sequence reads were trimmed (CutAdapt version 1.14), specifying a quality score of Q30, and then processed using CLC Genomics Workbench Version 11. Paired-end reads were merged, primer locations were trimmed, and processed reads were aligned to the SacCer3 reference genome. CLC low frequency variant caller was used to call variants, with required significance of 0.01%. Variant files were exported from CLC as VCF files and downstream analysis was performed in RStudio (version 1.2.1335), paired with custom python scripting. All sequence variants are provided in Table S22. Variants in wild-type, *rnr1D57N*, *rnr1Y285F, rnr1Y285F pGAL-RNR1*, *rnr1Y285A* and *rnr1Y285A pGAL- RNR1* were characterized previously (Lamb *et al*. 2021) and are used here to compare single and double mutants.

### Spearman Rank and Hierarchical Cluster Analysis

We used Spearman rank correlations among all genotypes (Fig. S2) to assess the relationship between genotype and mutation profile. Biological replicates from strains with lower mutation rates (e.g. wild-type, *msh3Δ*, *rnr1D57N*) were less well correlated than those from strains with higher mutation rates (e.g., *msh2Δ, msh6Δ, rnr1Y285A*), indicating that mutation events occurred more systematically (less stochastically) in these genetic backgrounds, consistent with specific mutation profiles. For example, the average correlation of all 7 wildtype biological replicates with all other samples was *r_s_* = 0.192, while the average correlation within the 7 wildtype biological replicates was *r_s_* = 0.191. In contrast, the *rnr1Y285A msh3Δ* biological replicates were much more highly correlated to one another (*r_s_* = 0.527) than they were with all other samples (*r_s_* = 0.264).

Hierarchical Cluster Analysis was performed on data that was grouped by biological replicates and the number of times the different classes of variants occurred within a genotype. Data was combined on the average of unique counts of variants from all the biological replicates within a genotype. The different classes of variants include 6 classes of single nucleotide variants (SNVs), single base A/T or G/C insertions and deletions, complex insertions and deletions, as well as mononucleotide variants (MNVs) and replacements (Replac.). MNVs are dinucleotide SNVs, where two neighboring nucleotides are both mutated, ex: CC> AT. Replacements are complex insertions or deletions, where the deleted or replaced base is a variant. Two examples include AAC > G and C > AT. Both MNVs and replacements are extremely low frequency events and rarely occur in our data set; neither had a significant impact on clustering.

The data was condensed based on genotype by combining biological replicates and adding the total number of times a variant was seen in the total number of replicates. This analysis does not take frequency into account and instead totals how many unique types of variants occur in a sample. If the same variant occurred in multiple biological replicates within a genotype it was counted as such.

Spearman correlation coefficients were calculated across all samples. Spearman correlation was chosen because we did not consistently observe linear relationships between variant frequencies in our data set, although Pearson correlation yielded nearly identical results. Correlation analysis was plotted in RStudio using ggplot visualization packages .

### Principal Components Analysis (PCA)

PCA was performed on the number of times a unique variant was observed within a genotype. PCA was plotted using the “factoextra” package in RStudio, with a singular value decomposition (SVD) approach using the prcomp() function.

### Determining SNV in trinucleotide context

A 3 bp window surrounding each *CAN1* SNV was identified. The average number of each SNV within this context, a total of 96 possible trinucleotide contexts (Lamb *et al*. 2021) was calculated for each genotype and divided by the number of times that context occurs in *CAN1*. (Alexandrov *et al*. 2015). The number of trinucleotide sequence contexts in *CAN1* was calculated using a sliding window approach utilizing python scripting. For each of the 96 different SNV changes in triplet context, the average number of SNVs in a genotype was divided by the number of times the triplet sequence context occurs in *CAN1*. This dataset was imported into R-studio and plotted via the barplot() function.

### Cluster Analysis for genotype-specific correlations

To identify variants specific to a particular genotype and to eliminate frequency bias from variants that occurred early in the growth of a particular sample, we condensed unique variants based on the number of biological replicates sequenced for that genotype. If a particular variant occurred in 4 out of 4 biological replicates it was represented as 1, if it occurred in 3 out of 6 replicates it was represented as 0.5. This gives an unbiased approach to score for the probability that a particular variant was present in a particular genotype, without considering the absolute frequency at which that variant occurred. These data were clustered on rows (or unique variants), after applying a row sum cutoff of greater than 2. This cutoff eliminates low frequency variants which are less likely to be driving the differences in mutation spectra that we observe. By clustering the data only on unique variants, it allows us to see groups of different *types* of variants in specific sequence contexts that are potentially diagnostic for a particular genotype.

To infer variants that were enriched in a particular genotype we divided the probabilities into four different bins (0-0.25, 0.26-0.50, 0.51-0.75, and 0.76-1.0). A variant was positively enriched in a genotype if it occurred at 0.76 or greater probability, and negatively enriched if it occurred below a 0.25 probability. Variants were grouped for motif enrichment (black boxes) based on the main branches in the dendrogram, paired with similar patches of enrichment on the heatmap. It is worth noting the majority of variants that were negatively enriched did not occur at all in a given genotype (light blue on heatmap). On average there was greater than a 2-fold increase in probability for variants that were positively enriched.

### CC Dinucleotide cluster analysis

A python script using a sliding window approach was used to identify all reference positions containing CC dinucleotides within *CAN1*. Our dataset was then subset to include only variants that occurred in these dinucleotide CC positions. Of the 138 CC dinucleotide contexts across *CAN1*, 110 (∼80%) were mutated, compared to 857/1,711 base pairs or ∼50% of the base pairs in *CAN1.* Unique CC run variants were clustered based on the number of times that variant occurred in each genotype, while accounting for (normalizing by) the number of biological replicates sequenced for each genotype, as described above. Heatmaps were plotted using the pheatmap package in RStudio and motif enrichment was performed using Berkeley web logos (Crooks *et al*. 2004).

### COSMIC SBS signature cluster analysis

We performed hierarchical clustering to compare the mutation spectra from our study with human mutation signatures, through an unbiased approach. The single base substitutions (SBS) COSMIC signatures from GRCh38 (v3.2- March 2021, https://cancer.sanger.ac.uk/signatures/downloads/) were combined with the normalized SNVs in trinucleotide context (Fig. 4) (Alexandrov *et al*. 2020). Hierarchical cluster analysis was then performed on these combined data as described above.

### Data Availability Statement

All strains and plasmids are available upon request. All *CAN1* sequences were uploaded to SRA BioProject PRJNA785873. Spatial Analysis of Functional Enrichment (SAFE) analysis can be found at: https://github.com/baryshnikova-lab/safepy). All mutation profile and SGA data are also available in the Supplemental tables. Representative code is available at: https://github.com/nalamb/CAN1-Paper.

## Results

### Pathways in DNA metabolism, including MMR, interact genetically with *rnr1* alleles

Previous work (Xu *et al*. 2008; Buckland *et al*. 2014; Watt *et al*. 2016) demonstrating the combined effect of *rnr1* alleles and *msh* deletions on mutation rates suggested that together these genes contribute to increased replication fidelity, and therefore they might show synergistic effects on cell fitness. We predicted that other pathways also interact genetically with *rnr1* alleles that altered dNTP pools. Therefore, we performed synthetic genetic array (SGA) analysis using each of three different *rnr1* alleles (Fig. 1 and Tables S3-S9). The *rnr1Y285A* query had the greatest number of genetic interactions, consistent with the Y285A mutation having the highest increase in dNTP levels.

**Figure 1.**
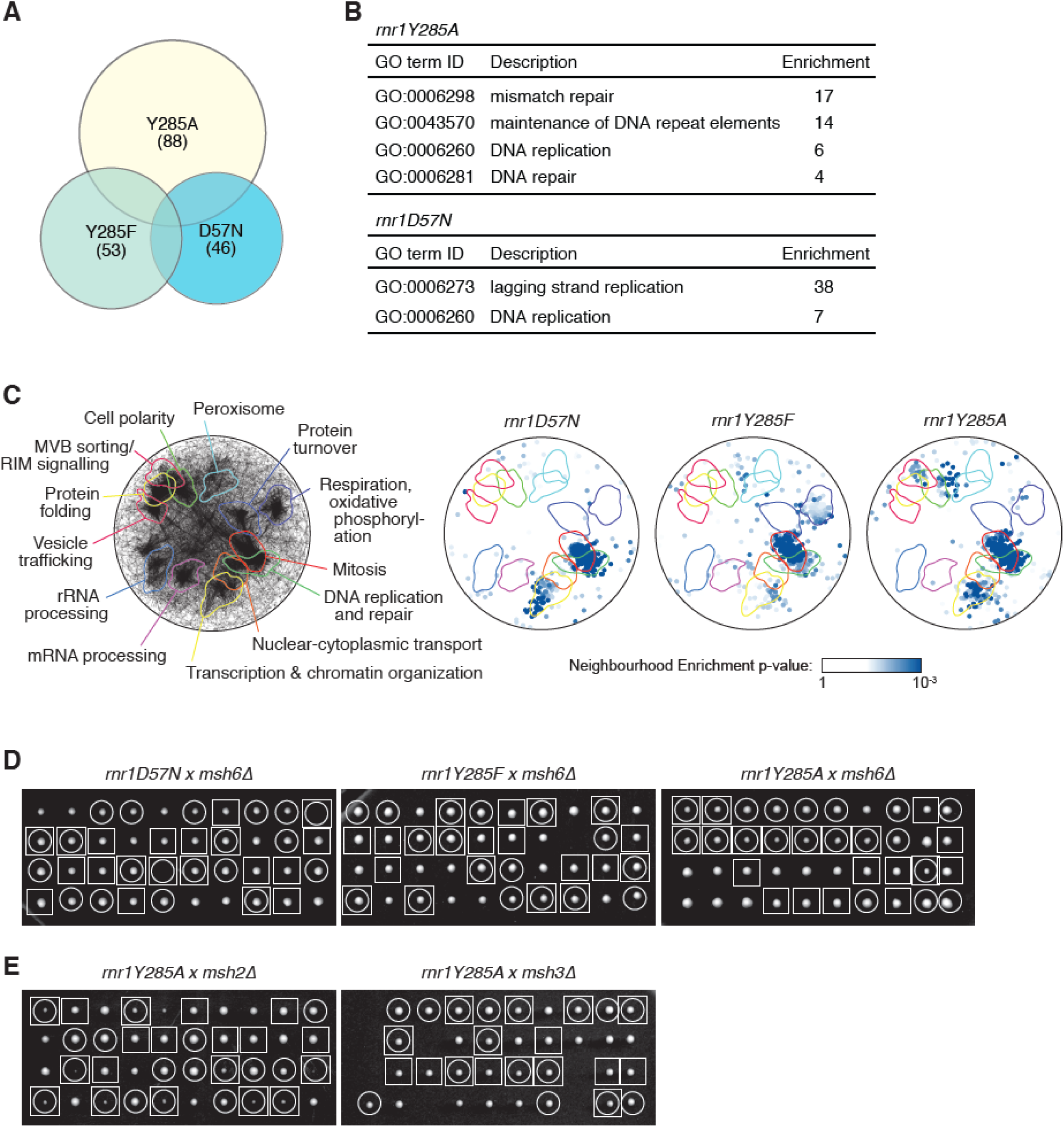
Genome-scale screens for synthetic fitness defects with *rnr1* alleles. (A) The overlap of the *rnr1* negative genetic interactions for the three SGA screens is plotted as a Venn diagram. The number of genes identified in each screen is indicated, as is the *rnr1* allele for each screen. (B) GO-term enrichments for the negative interacting genes from each *rnr1* screen are tabulated. The –fold enrichment for each term is indicated. Note that *rnr1Y285F* did not display any statistically supported enrichment. (C) Spatial analysis of functional enrichment. On the left, the yeast genetic interaction similarity network is annotated with GO biological process terms to identify major functional domains (Costanzo *et al*. 2016). 13 of the 17 domains are labeled and delineated by colored outlines. On the right, the network is annotated with negative genetic interactions from each *rnr1* SGA screen. The overlays indicate the functional domains annotated on the left. Only nodes with statistically supported enrichments (SAFE score > 0.08, p < 0.05) are colored. (D) Tetrad analysis of *rnr1* x *msh6*Δ crosses. Ten tetrads were dissected for each cross, and colonies were imaged after 3 days. Each column of 4 colonies is the 4 spores from a single meiotic ascus. Genotypes are indicated by circles (*msh6*Δ) and squares (*rnr1*). (E) Tetrad analysis of *rnr1Y285A* x *msh2*Δ and *rnr1Y285A* x *msh3*Δ crosses. Genotypes are indicated by circles (*msh2*Δ or *msh3*Δ) and squares (*rnr1Y285A*).

The three *rnr1* alleles showed surprisingly little overlap in their genetic interactions (Fig. 1A), supporting the idea that different dNTP levels and pool balances stress cells in different ways. Both the *rnr1Y285A* screen and the *rnr1D57N* screen showed enrichment for the GO term ‘DNA replication’ (Fig. 1B) in addition to displaying unique enrichments for ‘maintenance of DNA repeat elements’ and ‘DNA repair’ (*rnr1Y285A*) and ‘lagging strand replication’ (*rnr1D57N*). Despite having interactions with several DNA replication genes, the *rnr1Y285F* screen did not show any statistically supported GO term enrichment. To further assess the functional properties of each *rnr1* allele genetic interactions, we applied spatial analysis of functional enrichment (SAFE) (Baryshnikova 2016) to determine if any regions of the functional genetic interaction similarity yeast cell map (Costanzo *et al*. 2016) are over- represented for the negative genetic interaction gene sets (Fig. 1C). We found a statistically supported over-representation of the negative interacting genes in the DNA replication and repair neighborhood of the genetic interaction cell map for all three *rnr1* alleles, indicating that dNTP pool alterations impinge most dramatically on the DNA replication and DNA repair capacity of the cell. As with GO term enrichment, differences among the three *rnr1* alleles were also apparent in the SAFE analysis, with *rnr1Y285A* and *rnr1D57N* interactors being over-represented in the chromatin organization neighborhood compared with *rnr1Y285A*, and *rnr1Y285F* and *rnr1Y285A* interactors showing more over-representation in the mitosis neighborhood than did *rnr1D57N* interactors.

Most notably, we found that almost the entire MMR pathway (*MSH2*, *MSH3*, *MSH6*, *MLH1*, *PMS1* and *EXO1*) was identified specifically in the *rnr1Y285A* screen and the ‘mismatch repair’ GO term was very strongly and specifically enriched in the *rnr1Y285A* screen (Fig. 1C). This indicates a strong requirement for the mismatch repair pathway when dNTP pools are highly unbalanced. We validated the specificity of the genetic interaction between MMR and *rnr1Y285A* by performing tetrad analysis of *msh6*Δ crosses with each of the three *rnr1* alleles (Fig. 1D). Fitness defects in double mutant colonies were only evident in the *rnr1Y285A* cross. In tetrad analysis, *msh6*Δ, and particularly *msh2*Δ showed stronger fitness defects when combined with *rnr1Y285A* than did *msh3*Δ (Fig. 1E), consistent with the reduced viability observed in mutation rate experiments (Table 1). Our genetic interaction data indicate that all three *rnr1* alleles interface with DNA replication and repair pathways, and that *rnr1Y285A* might be expected to have a particularly dramatic effect in MMR deficient cells.

**Table 1.** Canavanine mutation rates.

| Table 1. Canavanine mutation rates |  |  |
| --- | --- | --- |
| Genotype | Mutation Rate [95% Confidence Intervals] | Fold Change |
| Wildtype (n=115) | $2.7 \times 10^{-7}$ [$2.3 \times 10^{-7} - 3.2 \times 10^{-7}$] | 1 |
| *msh2Δ* (n=24) | $6.7 \times 10^{-6}$ [$5.8 \times 10^{-6} - 7.8 \times 10^{-6}$] | 24.8 |
| *msh3Δ* (n=32) | $7.8 \times 10^{-7}$ [$6.1 \times 10^{-7} - 9.7 \times 10^{-7}$] | 2.9 |
| *msh6Δ* (n=24) | $1.8 \times 10^{-6}$ [$1.4 \times 10^{-6} - 2.2 \times 10^{-6}$] | 6.7 |
| *rnr1D57N* (n=91) | $7.3 \times 10^{-7}$ [$6.3 \times 10^{-7} - 8.3 \times 10^{-7}$] | 2.7 |
| *rnr1D57N msh2Δ* (n=24) | $2.0 \times 10^{-5}$ [$1.8 \times 10^{-5} - 2.3 \times 10^{-5}$] | 74.1 |
| *rnr1D57N msh3Δ* (n=24) | $2.2 \times 10^{-6}$ [$1.7 \times 10^{-6} - 2.7 \times 10^{-6}$] | 8.2 |
| *rnr1D57N msh6Δ* (n=24) | $5.6 \times 10^{-6}$ [$4.7 \times 10^{-6} - 6.6 \times 10^{-6}$] | 20.7 |
| *rnr1Y285F* (n=42) | $7.5 \times 10^{-7}$ [$5.8 \times 10^{-7} - 9.5 \times 10^{-7}$] | 2.8 |
| *rnr1Y285F msh2Δ* (n=24) | $2.9 \times 10^{-5}$ [$2.6 \times 10^{-5} - 3.2 \times 10^{-5}$] | 107.4 |
| *rnr1Y285F msh3Δ* (n=24) | $3.7 \times 10^{-6}$ [$3.0 \times 10^{-6} - 4.5 \times 10^{-6}$] | 13.7 |
| *rnr1Y285F msh6Δ* (n=24) | $2.3 \times 10^{-5}$ [$2.0 \times 10^{-5} - 2.6 \times 10^{-5}$] | 85.2 |
| *rnr1Y285F pGAL-RNR1* (n=24) | $7.8 \times 10^{-7}$ [$5.7 \times 10^{-7} - 1.0 \times 10^{-6}$] | 2.9 |
| *rnr1Y285F pGAL-RNR1 msh2Δ* (n=24) | $1.5 \times 10^{-5}$ [$1.3 \times 10^{-5} - 1.8 \times 10^{-5}$] | 55.6 |
| *rnr1Y285F pGAL-RNR1 msh6Δ* (n=12) | $1.6 \times 10^{-5}$ [$1.3 \times 10^{-5} - 1.9 \times 10^{-5}$] | 59.3 |
| *rnr1Y285A* (n=46) | $5.5 \times 10^{-6}$ [$4.7 \times 10^{-6} - 6.2 \times 10^{-6}$] | 20.4 |
| *rnr1Y285A msh2Δ* (n=33) | $2.5 \times 10^{-5}$ [$2.1 \times 10^{-5} - 3.0 \times 10^{-5}$] | 92.6 |
| *rnr1Y285A msh3Δ* (n=49) | $6.2 \times 10^{-5}$ [$5.7 \times 10^{-5} - 6.6 \times 10^{-5}$] | 229.6 |
| *rnr1Y285A msh6Δ* (n=33) | $1.0 \times 10^{-5}$ [$8.9 \times 10^{-6} - 1.2 \times 10^{-5}$] | 37.0 |
| *rnr1Y285A pGAL-RNR1* (n=22) | $1.5 \times 10^{-5}$ [$1.2 \times 10^{-5} - 1.7 \times 10^{-5}$] | 55.6 |
| *rnr1Y285A pGAL-RNR1 msh2Δ* (n=24) | $1.3 \times 10^{-5}$ [$1.1 \times 10^{-5} - 1.5 \times 10^{-5}$] | 48.2 |
| *rnr1Y285A pGAL-RNR1 msh3Δ* (n=36) | $7.3 \times 10^{-5}$ [$6.8 \times 10^{-5} - 7.9 \times 10^{-5}$] | 270.4 |
| *rnr1Y285A pGAL-RNR1 msh6Δ* (n=30) | $1.0 \times 10^{-5}$ [$8.7 \times 10^{-6} - 1.2 \times 10^{-5}$] | 37.0 |

### Combining *rnr1* and *mshΔ* alleles significantly increased mutation rates

The striking MMR interaction with *rnr1Y285A*, combined with the combinatorial effects of *rnr1D57N* and *msh2Δ* or *msh6Δ* (Xu *et al*. 2008), encouraged us to focus on the combined effects of *mshΔ* and *rnr1* alleles on mutation profiles. To determine whether the *rnr1 msh* double mutants displayed the anticipated increases in mutagenesis, we measured the forward mutation rates at *CAN1*.

The relative mutation rates for *msh* deletions were consistent with previous observations (Marsischky *et al*. 1996; Xu *et al*. 2008; Kumar *et al*. 2011b; Lang *et al*. 2013), with *msh2Δ* exhibiting the highest mutation rate, *msh6Δ* exhibiting a lower but still increased mutation rate and *msh3Δ* exhibiting a mild mutator effect in this assay (Table 2). The *rnr1* mutation rates increased with the increasing dNTP levels as expected (Kumar *et al*. 2010), with *rnr1Y285A* showing a 20-fold increase above the wildtype mutation rate. When *rnr1D57N* and *rnr1Y285F* were combined with *msh* deletions, the mutation rates exhibited the same hierarchy (*msh2Δ* > *msh6Δ* > *msh3Δ*), but with higher rates than either of the relevant single mutants. In most cases the mutation rate of the double mutant approximated the product of the single mutant mutation rates. The *rnr1Y285F msh6Δ* strain had a particularly large increase, 85-fold above wild-type, compared to the expected 19-fold given the mutation rates of the respective single mutants. Notably, *rnr1Y285A msh3Δ* exhibited a strong, synergistic mutator phenotype, indicating that the mutations generated in the presence of *rnr1Y285A* are substrates for Msh2-Msh3-mediated MMR. It is possible that the mutation rates for *rnr1Y285A msh2Δ* and *rnr1Y285A msh6Δ* are underestimates, due to fitness defects in these strains, consistent with our SGA results (Fig. 1) and (Watt *et al*. 2016).

**Table 2.** Total number of variants observed per genotype.

| Table 2. Total number of variants observed per genotype |  |
| --- | --- |
| Genotype | No. Variants |
| WT* | 267057 |
| *msh2Δ* | 120472 |
| *msh3*Δ | 122768 |
| *msh6*Δ | 95701 |
| *rnr1D57N* * | 205223 |
| *rnr1D57N msh2*Δ | 185107 |
| *rnr1D57N msh3*Δ | 72482 |
| *rnr1D57N msh6*Δ | 88642 |
| *rnr1Y285F* * | 171257 |
| *rnr1Y285F msh2*Δ | 94742 |
| *rnr1Y285F msh3*Δ | 79330 |
| *rnr1Y285F msh6*Δ | 104984 |
| *rnr1Y285F pGAL-RNR1* | 114080 |
| *rnr1Y285F pGAL-RNR1 msh2*Δ | 177905 |
| *rnr1Y285F pGAL-RNR1 msh6*Δ | 100200 |
| *rnr1Y285A* * | 106165 |
| *rnr1Y285A msh2*Δ | 153727 |
| *rnr1Y285A msh3*Δ | 98643 |
| *rnr1Y285A msh6*Δ | 166704 |
| *rnr1Y285A pGAL-RNR1* * | 176405 |
| *rnr1Y285A pGAL-RNR1 msh2*Δ | 110899 |
| *rnr1Y285A pGAL-RNR1 msh3*Δ | 97346 |
| *rnr1Y285A pGAL-RNR1 msh6*Δ | 101992 |

\*These variants were characterized in (LAMB *et al.* 2021)

Previous work characterized *rnr1Y285F* and *rnr1Y285A* (+/- *MSH2)* in backgrounds that carried a silenced *pGAL-RNR1* gene (i.e. grown in the absence of galactose) (Kumar *et al*. 2010; Buckland *et al*. 2014). We previously demonstrated that the mutation profiles of these strains were comparable to *rnr1Y285F* and *rnr1Y285A* without the *pGAL-RNR1* construct (Lamb *et al*. 2021). They are included here, with and without *MSH* genes to provide a comparison with that previous work. In general, *pGAL-RNR1* did not impact the observed mutation rates (Table 1) or profiles (Fig. S2, box2; Fig. S3) and were combined with the equivalent strains lacking *pGAL-RNR1* for some of our analyses.

### Distinct mutation profiles in cells lacking Msh2-Msh3- versus Msh2-Msh6-mediated MMR

Using our targeted deep sequencing approach, we defined and characterized mutation profiles in *rnr1 mshΔ* backgrounds, to derive mutation profiles from first principles (Lamb *et al*. 2021). First, we deleted *MSH2, MSH6* or *MSH3*, which encode the subunits of the MMR recognition MSH complexes (Tables S1, S2),(Lamb *et al*. 2021). We selected ∼2,000 canavanine resistant (Can^R^) colonies (selected samples) for each different genetic background, pooled them and extracted the pooled genomic DNA. These samples were subjected to paired-end deep sequencing of the *CAN1* gene. Selected samples exhibited an average variant frequency of 99.35% (Fig. S1A). We applied a permissive variant filter, based on the average frequency of individual variants in permissive samples (grown in the absence of selection) and the positions in which they occurred, to remove background noise resulting from PCR/sequencing errors (Lamb *et al*. 2021) (Fig. S1B). This approach identifies primarily base substitution and small insertion/deletion mutations; rearrangements will not be identified (Lamb *et al*. 2021). Further, there is a bias toward inactivating mutations, although we do observe about 3% synonymous mutations (data not shown). Nonetheless, the depth of sequencing and large number of total variants observed (Table 2) revealed novel insights into altered replication fidelity in the absence of either Msh2-Msh3- or Msh2-Msh6-mediated MMR.

Deletion of *MSH2* effectively eliminates all MMR activity and thus, retains errors made by replicative polymerases (Kunkel AND ERIE 2005). We also removed only Msh2-Msh6 (*msh6Δ*) or Msh2- Msh3 (*msh3Δ*) to learn more about substrate specificity and repair efficiency of each complex.(Sia *et al*. 1997; Harrington AND KOLODNER 2007; Xu *et al*. 2008; Kumar *et al*. 2011a; Lang *et al*. 2013; Buckland *et al*. 2014; Serero *et al*. 2014) Our work significantly increased the number of colonies sequenced and the sequencing coverage (Table 2). Previous studies focused primarily on *msh2Δ* and studies of *msh3Δ* and *msh6Δ* analyzed fewer than 100 colonies via Sanger sequencing (Sia *et al*. 1997; Harrington AND KOLODNER 2007; Xu *et al*. 2008), preventing statistically supported conclusions about variant types. In general, *msh2Δ, msh3Δ* and *msh6Δ* mutation profiles were consistent with previous studies performed in yeast (Sia *et al*. 1997; Harrington AND KOLODNER 2007; Xu *et al*. 2008; Kumar *et al*. 2011a; Lang *et al*. 2013; Buckland *et al*. 2014; Serero *et al*. 2014), but provided increased resolution to reveal novel specificities. Deleting *MSH2*resulted in a relative decrease in single nucleotide variants (SNVs) and increased deletions and insertions compared to wildtype, similar to previous observations (Lang *et al*. 2013; Serero *et al*. 2014). A/T single base deletions dominated the mutation profile in *msh2Δ* (Table S10, Fig. 2B, C, D). In contrast, the *msh6Δ* spectrum was dominated by SNVs, which represented >80% of the mutations (Table S10, Fig. 2A, B), although the *types* of SNVs generated were similar for *msh2Δ* and *msh6Δ* (Table S10, Fig. 2B). In *msh3Δ* cells there was a similar proportion of deletion events compared to *msh2Δ*, but the types of mutation varied. There was a marked increase in G/C -1 bp deletions and complex deletions and insertions (> 1 base pair) compared to wild-type, *msh2Δ* or *msh6Δ*, consistent with the preference of Msh2-Msh3 for binding larger IDLs (Habraken *et al*. 1996; Surtees AND ALANI 2006). Approximately 30% of the mutations that accumulated in *msh3Δ* were SNVs, but again the distribution was distinct, with increased CG>GC and TA>AT changes, despite the fact that these errors should be efficiently repaired by Msh2-Msh6 (Genschel *et al*. 1998; Bowers *et al*. 1999); TA>AT errors in *msh3Δ* were more frequent than observed in wildtype (Table S10, Fig. 2A). There were fewer TA>CG changes in *msh3Δ* compared to *msh2Δ* and *msh6Δ* (Table S10, Fig. 2A). These observations are consistent with Msh2-Msh3 also playing a role in correcting a specific subset of misincorporation events (Harrington AND KOLODNER 2007; Kumar *et al*. 2011a).

**Figure 2.**
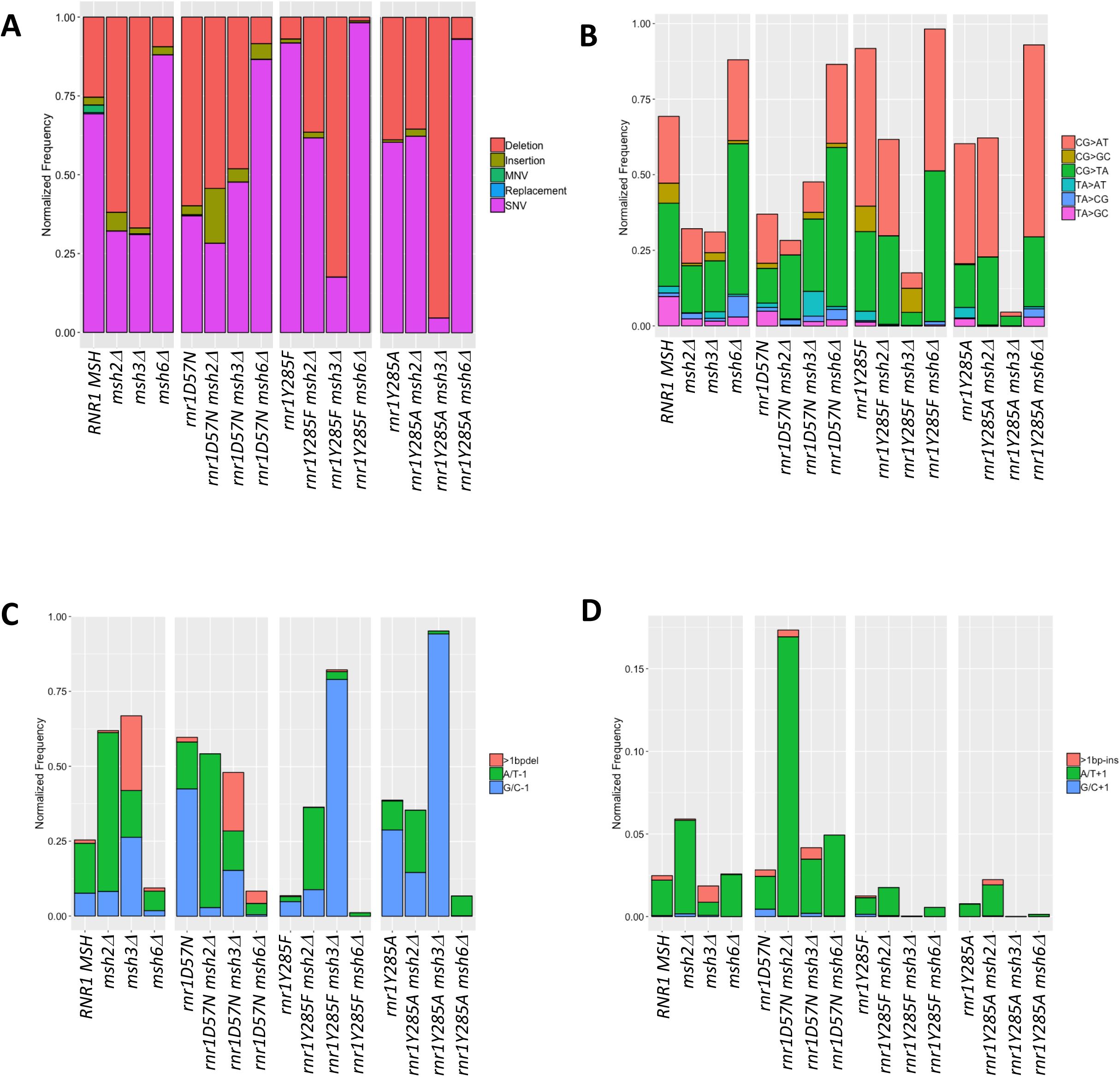
Mutation Spectra are distinct across genotypes. (A) The relative distribution of SNVs, deletions, insertions, MNVs and replacements by genotype. (B) The SNV spectra normalized relative total variants. (C) The deletion spectra normalized relative to total variants. (D) The insertion spectra normalized relative to total variants.

### Deep sequencing reveals systematically genotype-specific mutation spectra in *rnr1 mshΔ* genetic backgrounds

In assessing the ability of Msh2-Msh3 and Msh2-Msh6 to direct repair of replication errors using *msh* alleles, we are limited to evaluating repair of those mutations that arise because of replication error. Due to the inherent proofreading activity of Polδ and Polε, this rate is low and certain mutations are very rare (Kunkel 2009; Arana AND KUNKEL 2010; Kunkel 2011). However, *rnr1* alleles generate distinct mutation profiles, including variants that are rare in wild-type backgrounds (Kumar *et al*. 2011b; Buckland *et al*. 2014; Watt *et al*. 2016; Lamb *et al*. 2021). To evaluate the specificity of MMR MSH complexes for a broader range of mutations and to assess the impact of combined genotypes on mutation profiles, we combined *msh* deletions with *rnr1* alleles. (Lamb *et al*. 2021) Notably, dNTP pools are likely elevated in cancer cells (Aye *et al*. 2015; Mathews 2015), including those caused by defects in MMR. Therefore, the effect of the combination of *mshΔ* and altered dNTPs on replication fidelity could result in unique mutation signatures observed in tumors.

We performed targeted deep sequencing of *rnr1 mshΔ* mutants and characterized mutation events in two ways (Table S10), as previously described (Lamb *et al*. 2021). First, we determined the number of a specific variant type, i.e., the number of C>A changes, at different positions along *can1* (“unique counts”) (Counts in Table S10). Second, we calculated the frequency at which each of these unique variants occurred, i.e., the combined frequency of all C>A changes at any position along *can1* (“sum of frequencies”) (Freq. in Table S10). These analyses allowed us to determine whether different *types* of mutations occurred at specific locations in a genotype-dependent manner, independent of frequency, and whether variant frequencies were altered in a significant way by genotype (Counts/Freq. in Table S10). A decreased number for “unique counts” combined with unchanged or increased “sum of frequencies” would indicate that variant type is more localized, possibly indicating a mutational hotspot. For instance, *msh6Δ* exhibited the highest proportion of unique events contributing to the mutation spectrum (Counts/Freq. = 1.62; Table S10). In contrast, *rnr1Y285F msh3Δ* and *rnr1Y285A msh3Δ* exhibited the lowest proportion of unique variants; the mutation spectra were instead dominated by G/C single base deletions, which occur at high frequencies (Table S10).

We used Spearman rank correlation and cluster analysis to determine the quantitative relationship between mutation profile and genotype (see Materials and Methods). All unique variants for all genotypes were assessed in parallel, based on both the presence and frequency of unique variants, as described above. In general, biological replicates of the same genotype clustered because their mutational profiles were highly correlated (Fig. S2, Table S11). Therefore, we combined variants from all biological replicates within a genotype for the remainder of our analyses. Hierarchical cluster analysis using Spearman rank correlations based on the profile of unique variants between genotypes was consistent with genotype-specific mutation profiles (Fig. 3A, Fig. S2), as was principal component analysis (PCA) based on unique variants (Fig. 3B). Combined, these results indicated that it is possible to distinguish among genotypes based on unique variant profiles observed from *can1* deep sequencing. Below, we parse these trends to develop genotype-specific mutation profiles.

**Figure 3.**
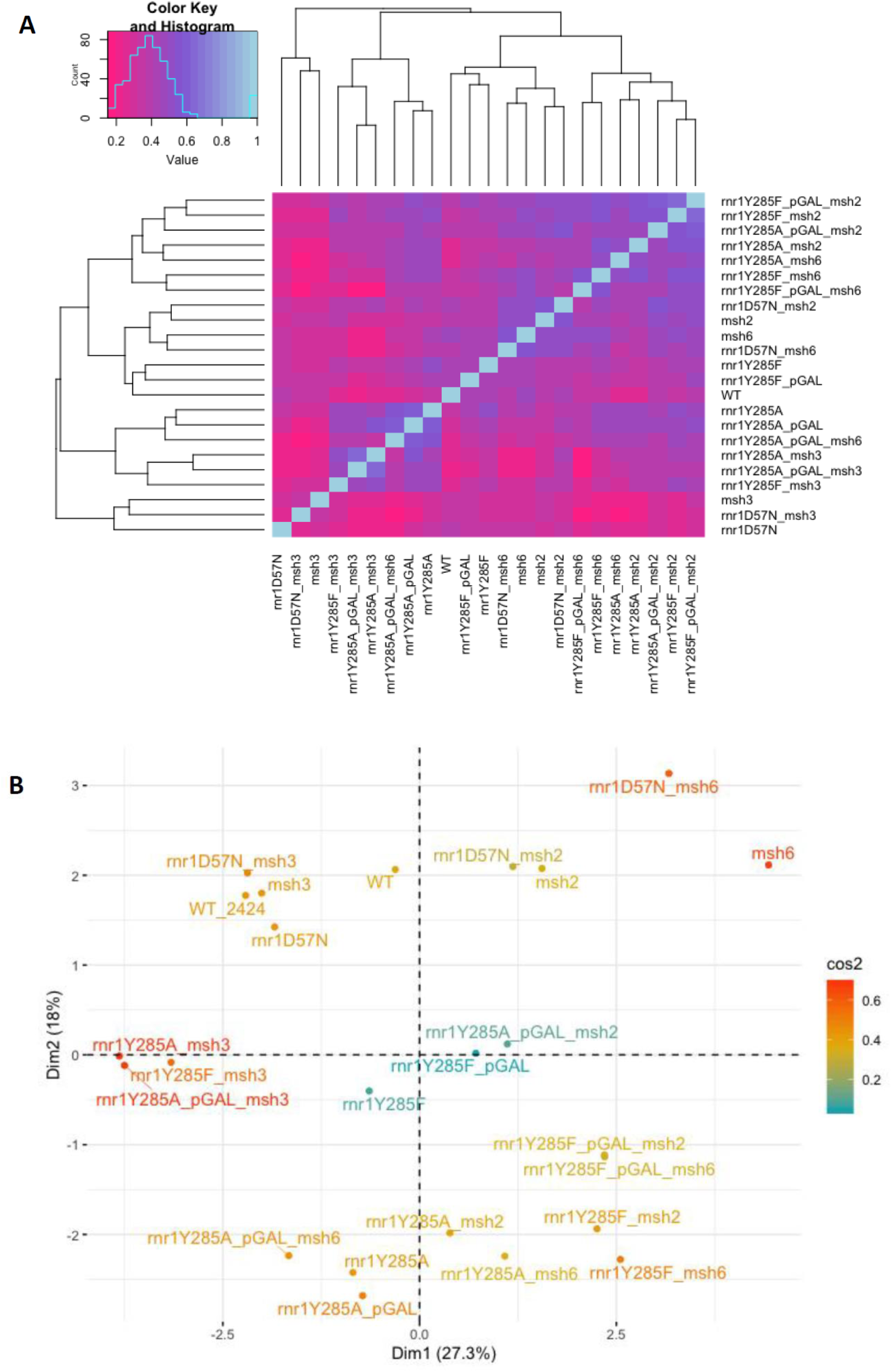
Distinct genotypes share unique features. (A) Hierarchical cluster analysis using Spearman Rank correlation on all the distinct genotypes in our study. Data was clustered based on the unique counts of the 14 different classes of variants that occurred in each genotype. The histogram shows the distribution of correlation coefficients across samples. (B) Principal components analysis performed on all unique variants from biological replicates within a genotype.

**Figure 4.**
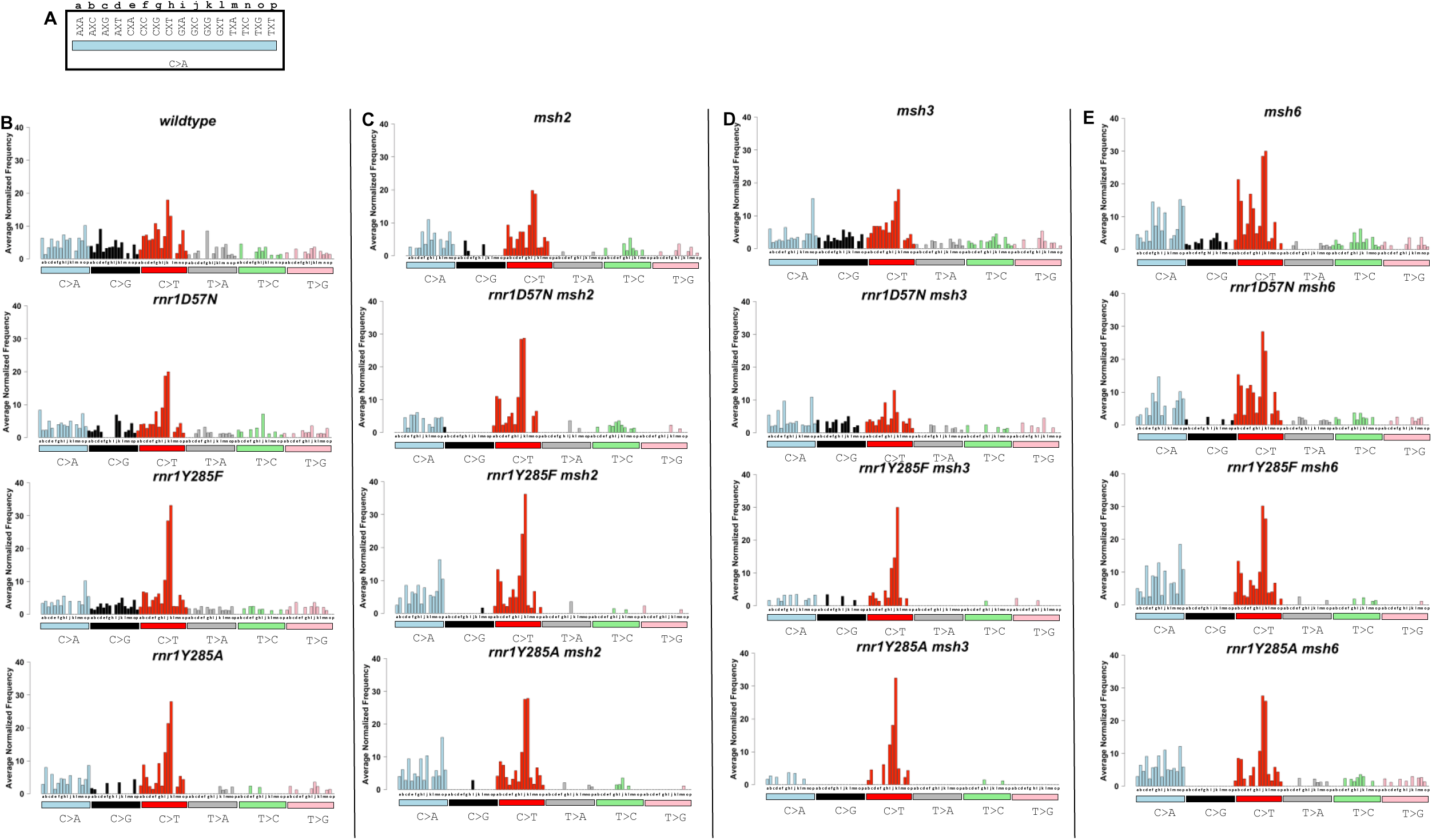
The average number of each SNV as it occurs in unique triplet nucleotide context differs by genotype. Bars are colored according to the six different types of SNVs. (A) The 16 different triplet contexts are lettered for display purposes. The variant change (C>A, turquoise bar) occurs at the middle nucleotide marked X in each triplet context. The same triplet context is repeated for each possible variant in the following panels. (B) Wildtype and single *rnr1* alleles. (C) Genotypes with *MSH2* deleted (D) Genotypes with *MSH3* deleted. (E) Genotypes with *MSH6* deleted.

### *rnr1* alleles combined with MMR deletion result in mutations within distinct sequence contexts and motifs

When *msh2Δ* was combined with each *rnr1* allele, distinct mutation spectra (i.e. the proportion of each type of variant) were observed, revealing mechanistic insights into MMR and replication fidelity (Table S10, Fig. 2A). The *rnr1D57N*, *rnr1Y285F* and *rnr1Y285A* profiles exhibited an increase in G/C-1 deletions relative to A/T-1 deletions compared to wildtype. The opposite trend was observed in increases the proportion of G/C-1 variants, A/T-1 deletions are preferentially repaired in the presence of MMR. There was also a significant increase in A/T+1 insertions in *rnr1D57N msh2Δ* relative to either single mutant. Finally, while the *rnr1Y285F* and *rnr1Y285A* SNV profiles were biased toward CG>TA and CG>AT SNVs, *rnr1Y285F msh2Δ* and *rnr1Y285A msh2Δ* were almost completely dominated by these variants, with proportions that differed from either single mutant (Table S10, Fig. 2). These SNV profiles indicated that: 1) DNA polymerases primarily generated these errors when dCTP and dTTP were modestly skewed and elevated (*rnr1Y285F*) and 2) these elevated frequencies began to saturate MMR activity and/or these errors are inefficiently repaired by MMR.

Deleting *MSH6* in combination with *rnr1* alleles, so that only Msh2-Msh3-mediated MMR was present, also resulted in unique shifts in mutagenesis across *can1* (Table S3, Fig. 2A). The effect was most dramatic in *rnr1Y285F msh6Δ* with a profile that was almost completely dominated by CG>TA and CG>AT SNV errors (Table S10, Fig. 2B, C), indicating that Msh2-Msh3 is not efficient in correcting these mismatches. The proportion of CG>AT transversions was even higher in *rnr1Y285A msh6Δ*, although most of the variant classes were still observed. In both *rnr1Y285F msh6Δ* and *rnr1Y285A msh6Δ* there was almost a complete loss of the G/C -1 bp deletions that were observed at an increased frequency in *rnr1Y285F* and *rnr1Y285A* (Table S10, Fig. 2C,D), consistent with efficient repair of G/C -1 bp slippage events by Msh2-Msh3-mediated MMR. Strikingly, the opposite was observed in *msh3Δ* genotypes. In both *rnr1Y285F msh3Δ* and *rnr1Y285A msh3Δ*, there was a dramatic increase in G/C -1 bp deletions compared to either single mutant, almost to the exclusion of other variants, indicating that Msh2- Msh6 was unable to correct this type of error.

### Genotype-specific susceptibility to mutation at specific positions within *CAN1*

The mutation profiles described above indicated the possibility of genotype-specific positions within *CAN1* that were susceptible to mutation (Table S12-S17). We identified and defined three classes of positions vulnerable to mutation: 1) susceptible positions were those in which the same variant was observed at the same position in >50% of biological replicates, 2) increased susceptibility sites, which also had a variant frequency above wild-type and 3) highly susceptible positions which exhibited a variant frequency >2-fold above wildtype. All variants were more likely to occur at or adjacent to repetitive sequences, although the specific repetitive sequences varied by genotype. There were more susceptible positions, with higher mean frequencies, in *msh2Δ* (49 positions) and *msh6Δ* (96 positions) than in *msh3Δ* (18 positions). Of the 49 susceptible positions in *msh2Δ*, 9 were unique to this genotype. The remaining positions overlapped with those observed in *msh3Δ* and/or *msh6Δ*. Approximately 2/3 (61) of the susceptible mutations in *msh6Δ* were specific to this genotype. None of the *msh3D* susceptible sites were unique to *msh3Δ*; 4 were also observed in *msh2Δ* while the remaining 9 were also observe in *msh6Δ*, albeit at different frequencies. The highly susceptible *CAN1* positions that we observed in *msh2Δ* overlapped previously observed “hotspots” (Buckland *et al*. 2014; Watt *et al*. 2016); we identified additional susceptible positions, particularly in repetitive sequences.

We also characterized novel increased susceptibility sites in *msh6Δ* and *msh3Δ*. The *msh6Δ* exhibited 13 highly susceptible positions, all SNVs, which were not observed in wildtype and occurred almost exclusively in repetitive sequence contexts, including at dinucleotides and in runs 3bp or greater in length. In *msh2Δ*, 11 highly susceptible positions were observed, mostly deletions within repetitive runs, with few positions overlapping with *msh6Δ.* We previously identified novel susceptible positions in *rnr1D57N*, *rnr1Y285F* and *rnr1Y285A* (Lamb *et al*. 2021), many of which were associated with repetitive DNA sequences, particularly insertions and deletions.

We observed much more distinct profiles of susceptible positions in *CAN1* when *msh* deletions were combined with *rnr1* alleles, typically differing from the individual single mutants in the number of replicates affected and/or the frequency of mutation. The susceptible positions in *rnr1Y285A msh2Δ* identified in our study largely overlapped with previously identified “hotspots” (Kumar *et al*. 2011b; Buckland *et al*. 2014), but also revealed new susceptible positions. One noteworthy position is 32,940, which occurs in A-rich sequence (CCAA<u>G</u>AAAA) and is susceptible to G>T mutation in *msh2Δ rnr1Y285A* and *msh2Δ rnr1Y285A-pGAL*. The variant frequency at this position increased synergistically in *rnr1Y285A-pGAL msh2Δ* and *rnr1Y285A msh2Δ* double mutants, occurring at a frequency at least 10- fold greater than most single mutants. G>A SNVs also occurred in a variety of *msh6Δ* genotypes at this position, indicating decreased replication fidelity in this context. Notably, the majority of susceptible positions in *rnr1Y285A msh2Δ* are also susceptible in *rnr1Y285A* but not *msh2Δ*, indicating that highly skewed/elevated dCTP and dTTP levels promoted specific errors to a level where MMR was invoked, and approached saturation of the MMR system. By contrast, *rnr1Y285A msh6Δ* and *rnr1Y285A msh3Δ* tended to exhibit susceptible positions that were distinct from either single mutant, consistent with different specificities of Msh2-Msh3 and Msh2-Msh6-directed repair in the presence of a distinct mutational baseline observed in the presence of *rnr1* alleles.

### Insertions and deletions occur in distinct sequence contexts depending on genotype

As noted above, susceptible *CAN1* positions tended to be in or near repetitive sequences. We also specifically noted increased insertion and deletion events in repetitive runs, similar to previous work (Lujan *et al*. 2014; ST Charles *et al*. 2015). Within *CAN1*, there are only short homopolymer runs and A/T repeats are more abundant than G/C repeats: 266 AA/TT dinucleotides versus 160 GG/CC dinucleotides; 98 mononucleotide runs of 2 or more A/T bases versus only 32 G/C runs. These ratios are consistent with the proportion of A/T versus G/C deletions we observed in wildtype (Fig. 2C & 1D). 96.1% of all A/T -1 bp deletions occurred in a repetitive run of 3 bp or greater. 89.6% of G/C deletions occurred in a repetitive run of 2 or more nucleotides. Notably, some G/C deletions occurred adjacent to repetitive runs in a genotype-specific manner. For example, a GΔ at position 31971 occurred in 4 out of 7 *rnr1D57N* biological replicates, between an A dinucleotide and C dinucleotide (AT<u>AA</u>G<u>CCAA</u>). CΔ deletions at positions 32398 (G<u>AAA</u>CGTAG) and 32630 (<u>CCTT</u>CG<u>TTTT</u>) were specific to r*nr1Y285A*. A GΔ at position 33018 was specific to *msh3Δ* ; it is not flanked by a repetitive run but dinucleotides are nearby upstream and downstream (<u>GG</u>ATGT<u>AA</u>C).

Complex insertions and deletions (i.e. involving more than one nucleotide) were rare, but occurred at increased frequency in *msh3Δ* . The majority of these complex events, especially insertions, were observed in a single biological replicate. The complex insertion of CT in the repetitive run stretching from positions 33206- 33215 (CTTAAG<u>CT</u>CTCTC) is noteworthy. It was observed almost exclusively in *msh2Δ* genotypes, and more frequently when paired with *rnr1* alleles. The increased CT insertion in *rnr1 msh2*Δ genotypes indicates that positions 33206-33215 were particularly susceptible to mutation when dNTPs were elevated, even by a small amount as is the case in *rnr1D57N*. However, the CT insertion was very efficiently repaired by MMR, via either Msh2-Msh3 or Msh2-Msh6 directed repair as it was not observed in either *msh3Δ* or *msh6Δ*.

### Unique mutation signatures revealed by analysis of SNV trinucleotide context

To gain mechanistic information about sequence context that might influence either nucleotide misincorporation events or MMR, we determined the average number of times a SNV was observed in a particular triplet context per genotype, normalized by the number of times the triplet context occurs in *CAN1* (Fig. 4). Notably, C→T changes (red bars, Fig. 4), particularly in GCC and GCG sequence contexts, dominated in all genotypes, but most dramatically in *rnr1Y285F* and *rnr1Y285A* samples, as we previously observed (Lamb *et al*. 2021) (*r_s_* = 0.7284; Table S18). Whole genome sequencing also previously detected the prevalence of mutations in the GCC and GCG sequence contexts are mutated in *rnr1Y285A* (Watt *et al*. 2016), possibly as a result of increased extension after misincorporation of dTTP, which is in excess, superseding proofreading (Kunkel AND SONI 1988).

The wildtype SNV trinucleotide pattern was highly correlated with that of *msh3Δ* (*r_s_* = 0.7351), *rnr1D57N msh3Δ* (*r_s_* = 0.6752) and *rnr1D57N* (*r_s_* = 0.6682). No C>T SNVs occurred in GCT context in these four genotypes, while they were observed in all *rnr1Y285A* and *rnr1Y285F* double mutant backgrounds, albeit at relatively low frequencies. This example shows an error that was specific to skewed increases in dCTP and dTTP and was efficiently repaired by Msh2-Msh6.

We also observed a decrease in the number of unique SNVs that occurred in trinucleotide context when *rnr1Y285F* or *rnr1Y285A* alleles were paired with *msh2Δ, msh3Δ,* or *msh6Δ*. Calculation of Spearman correlation coefficients of each SNV in unique trinucleotide context revealed that these genotypes are also highly correlated to one another, with *rnr1Y285F/A msh6Δ* or *msh2Δ* double mutants showing the highest correlation values (Table S18). We observed a complete loss of C>G SNVs in all trinucleotide contexts in *rnr1Y285A msh3Δ*, *rnr1Y285A msh6Δ*, and *rnr1Y285F msh6Δ* backgrounds, as noted above (Fig. 2, Table S10). The C>G variant rarely occurred in *msh2Δ* backgrounds, indicating that the replicative polymerases rarely generate these errors. It is noteworthy that double mutant profiles often showed distinct changes in the SNV signatures, compared to either single mutant. For example, *rnr1Y285A* sustained several C>G mutations in multiple trinucleotide contexts, which were also observed in *rnr1Y285A msh2Δ,* but these were completely absent in *rnr1Y285A msh3Δ* and *rnr1Y285A msh6Δ*. Thus, there were combinatorial effects on the mutation spectra of specific MMR deletions in the presence of *rnr1* alleles. Different *rnr1* alleles paired with different MMR deletions result in distinct and unique mutational fingerprints.

### Novel error substrate specificities revealed for Msh2-Msh3 and Msh2-Msh6 complexes

All the analysis of variant profiles and position effects pointed to distinct genotype-specific differences in mutation and context when *MSH3* versus *MSH6* was deleted. Using all single and double mutant variant data, we performed hierarchical cluster analysis of all unique errors to identify larger sequence contexts driving distinct mutational signatures (Figs. 5). Genotypes were clustered based on the types of variants that were differentially enriched as a function of genotype, using Pearson correlation (Spearman correlation analysis yielded similar results) (Fig. 5). It is worth noting that overall, clusters were similar to those observed in Fig. 3, based on variant frequencies, which suggests that unique variants were the main drivers of genotype-specific mutation profiles. We identified differentially enriched variants and performed motif analysis (12 base window) to determine whether broader sequence context influenced the occurrence of these variants, indicating the same biological mechanism. We identified several motifs that were positively or negatively enriched in different genetic backgrounds. MMR status appeared to be the primary driver for enrichment, with G/C-1 variants positively enriched within G/C homopolymeric runs in *msh3Δ* genotypes, A/T-1 variants within A/T runs and several SNVs were positively enriched in *msh2Δ* and *msh6Δ* genotypes (see examples in Fig. 5).

**Figure 5.**
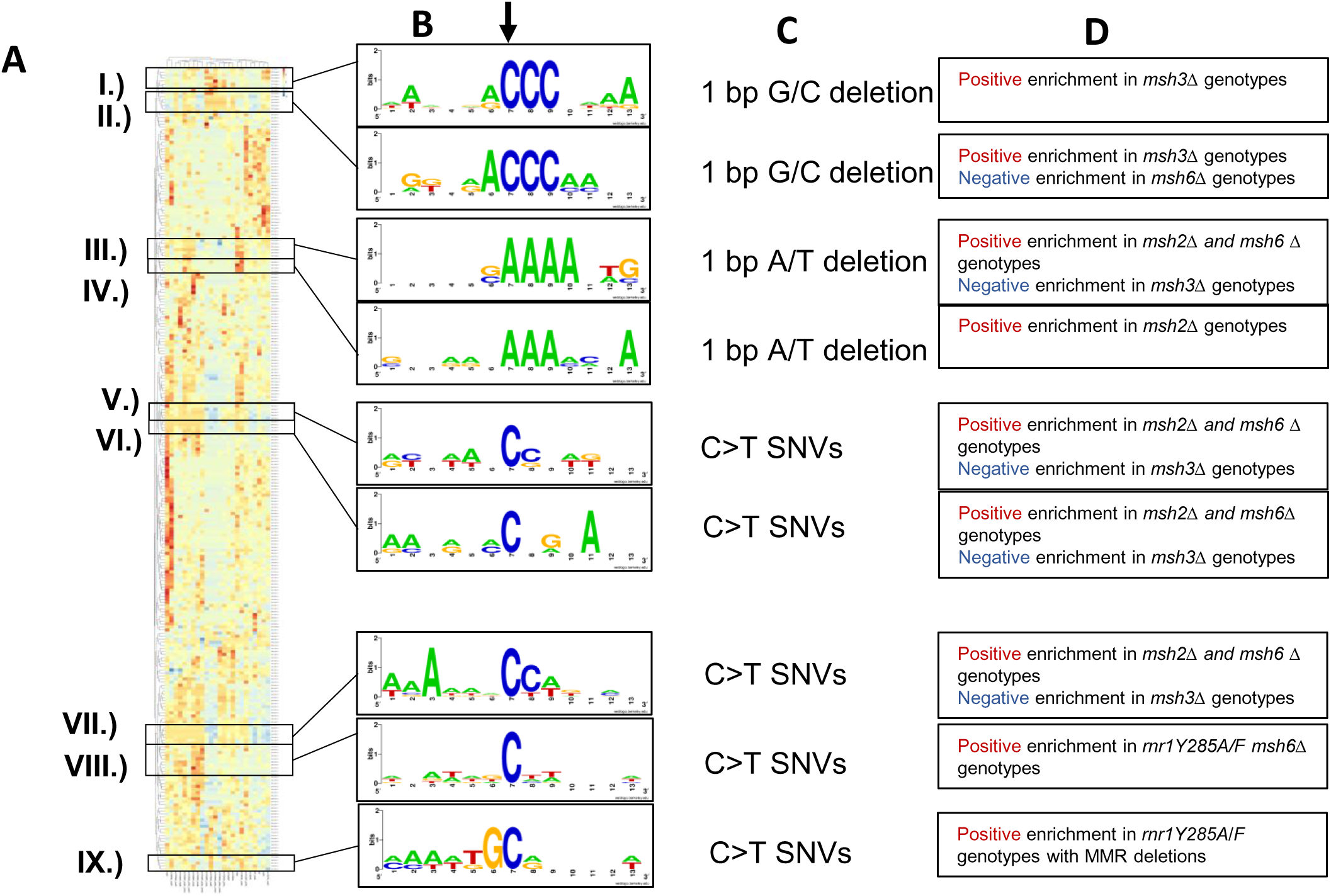
Variants that occur in unique sequence contexts cluster together Hierarchical cluster analysis of all unique variants within our study by genotype (A) A heatmap displaying Pearson correlation enrichment value for a given variant between genotypes, with notable clusters boxed in black. (B) 12 base window motif enrichment on sequence contexts surrounding the notable clusters. (C) The type of variant observed in the center of sequence context from B. (D) Summary of genotypes which show negative or positive correlation in each cluster.

Some of the clusters with the most pronounced differential enrichment between genotypes were SNVs that occurred in CC dinucleotide context (Fig. 6). As observed previously (Watt *et al*. 2016; Lamb *et al*. 2021), CC/GG dinucleotides bordered by G on either the 5’ or the 3’ side were frequently mutated in the presence of *rnr1* alleles, particularly *rnr1Y285F* and *rnr1D57N*. This was not observed with *msh* alleles. MMR status, (especially *msh3Δ* or *msh6Δ*) in combination with these *rnr1* alleles appeared to reverse this bias with negative enrichment. The majority of SNVs that occurred in other CC dinucleotide contexts were positively enriched in *msh2Δ* and *msh6Δ* samples, but negatively enriched in *msh3Δ* (Fig. 6 II, III, & IV), indicating that Msh2-Msh6 was uniquely required to recognize and direct repair of misincorporation events in the CC dinucleotide context.

**Figure 6.**
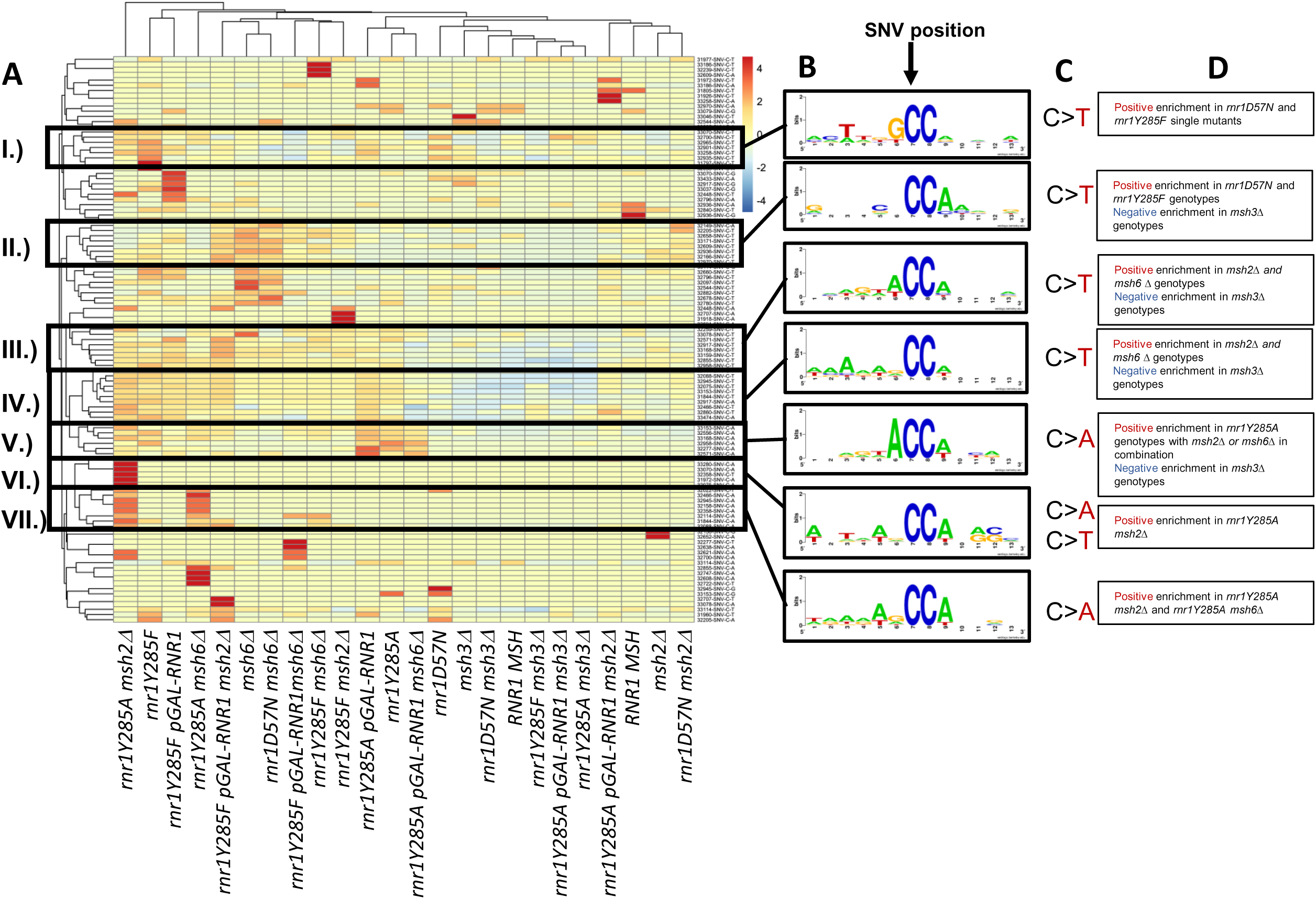
SNVs occur in C-C dinucleotide sequence contexts with differential enrichment between genotypes. (A) Hierarchical cluster analysis of all SNVs that occur at CC dinucleotides. Clusters of interest are boxed in black, labeled by roman numerals. (B) Motif enrichment of a 12 base window surrounding the mutated nucleotide was performed using Berkley web logos. The mutated base is at the 7^th^ nucleotide position in all logos, indicated by the black arrow. (C) The most predominant type(s) of SNV in the cluster are displayed. (D) A summary of genotypes that show negative or positive correlation in each cluster.

Motif enrichment analysis also revealed synergistic increases in G/C-1 variants within specific sequence contexts when *rnr1Y285F/*A and *msh3Δ* alleles were combined (Fig. 7). We previously found that *rnr1Y285F* and *rnr1Y285A* single mutants showed an increase in G/C single base deletions (Lamb *et al*. 2021). Here we found that *msh3Δ* does as well (Fig. 2, Table S10). Individual positions that sustained G/C errors along *CAN1* were often *msh3Δ-*specific (Table S10) or *rnr1Y285F/A*-specific (Lamb *et al*. 2021). Notably, the frequency of G/C-1 variants within sequence contexts that contain G or C runs increased synergistically in the *rnr1Y285F/A msh3Δ* double mutants (Fig. 7). In contrast, G/C-1 variant frequency was neutral or negatively enriched in all other genotypes, consistent with apparent specificity of Msh2-Msh3 for directing repair of G/C single base deletions (Fig. 2, Table S10, Fig. 5). Loss of Msh2-Msh3 resulted in increased G/C single base deletions in homopolymer runs bordered by G/C rich sequence on the 5’ side of the run (Fig. 7A-C). This error in this context occurred rarely in *rnr1Y285F/A* alone. There was a significant increase in G/C-1 mutations in *rnr1Y285F msh3Δ* double mutants. In contrast, G/C runs bordered by A/T nucleotides were more prone to mutagenesis in *rnr1Y285F/A* than in *msh3Δ* single mutants (Fig. 7D-F). The frequency of these variants directly bordered by A/T increased synergistically when *MSH3* was deleted in the presence of *rnr1Y295F/A*, but not when *MSH6* was deleted, indicating Msh2-Msh3 has specificity in directing repair of G/C -1 deletions in repetitive G/C context.

**Figure 7.**
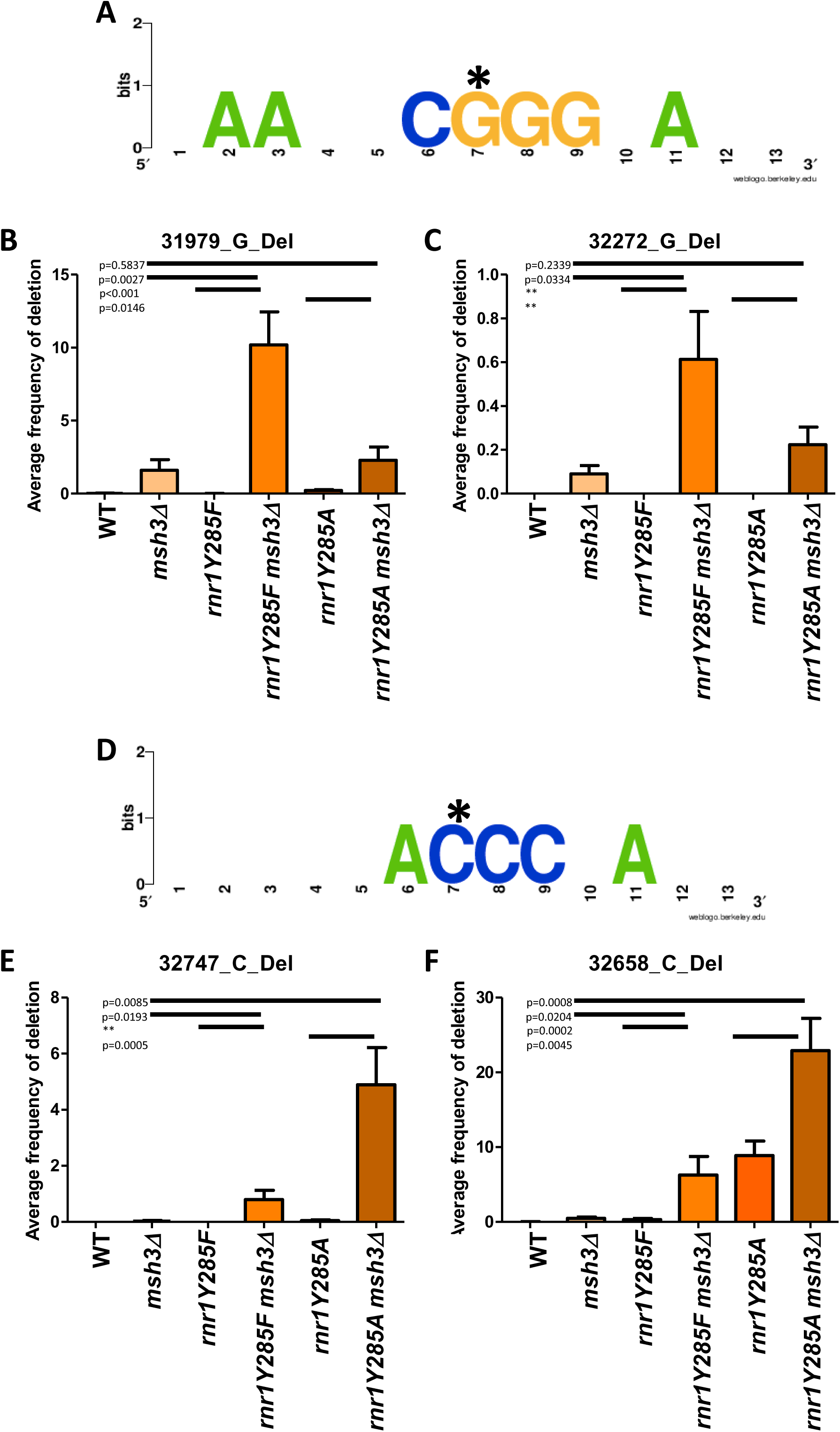
G/C single base deletions that show synergistic increases in variant frequency in *rnr1Y285A msh3Δ* genotypes. (A) Motif enrichment of a 12 base window surrounding two G deletions (starred nucleotide) that are specific to genotypes with *msh3Δ.* The asterisk indicates the deleted base. (B) The average variant frequencies from biological replicates in wildtype, *rnr1Y285F/A* and *msh3Δ* genotypes for the single base deletion that occurred at position 31979 is plotted. (C) The average variant frequencies across replicates for the G deletion at 32272. In both cases (B and C) there are very few events in the single mutants, but significant frequencies in double mutant backgrounds. (D) Motif enrichment for two C deletions that are specific to genotypes with *rnr1Y285F* and *rnr1Y285A*, but increase synergistically in double mutants with *msh3Δ* in combination. The asterisk indicates the deleted base. (E) The average variant frequencies across replicates for the G deletion at 32747. (F) The average variant frequencies across replicates for the G deletion at 32658. In E and F, this event occurs in *rnr1Y285A*, but not *rnr1Y285F* or *msh3Δ.* It occurs at increased frequencies in double mutant backgrounds. Error bars indicate the standard error of the mean for frequencies. The p values were generated by t-tests comparing average frequencies. A double asterisk indicates that a t-test was not possible because one of the genotypes had an average frequency of 0.

## Discussion

Utilizing a *CAN1* selection-based deep sequencing approach (LAMB *et al*.), we characterized mutation spectra in *mshΔ* and *rnr1 mshΔ* double mutant genotypes. While we likely missed non-inactivating mutations and more significant rearrangements that might occur, the sequencing depth afforded by our approach allowed us to expand our understanding of mismatch repair substrate recognition as well as the combined effects of MMR defects and altered dNTP pools on replication fidelity. By using *rnr1* backgrounds that alter the *type* and *frequency* of mutations sustained, we revealed previously unrecognized specificities for the MMR initiation complexes, Msh2-Msh3 and Msh2-Msh6. The combinatorial effects that we find highlight the importance of studying mutation signatures in different genetic contexts.

### Different mechanisms of mutagenesis result from distinct elevations in dNTP levels

In *rnr1D57N msh2Δ* the mutation rate increased 74-fold above wildtype and 3-fold above *msh2Δ*, yet the mutation spectrum of *rnr1D57N msh2Δ* is closely related to *msh2Δ*, with the exception of an increase in A/T+1 insertions in the double mutant (Fig. 2). The same is true of *rnr1D57N msh3Δ* and *rnr1D57N msh6Δ;* their mutation spectra are most closely related to those of *msh3Δ* and *msh6Δ*, respectively, despite high increases in mutation rates (Fig. 2, Tables 1 & S3). Therefore, the elevated dNTP pools in *rnr1D57N*, which resulted in a mutation spectrum similar to wildtype (LAMB *et al*.), with the notable exception of G/C-1 deletions, did not substantially drive the type of mutation generated. The low frequency variants that accumulate in *rnr1D57N* were effectively repaired by MMR, in general, and even the absence of MMR did not result in an overt fitness defect in *rnr1D57N* (Fig. 2, Table S7, S12). We conclude that the balanced dNTP increases in *rnr1D57N* alter mutagenesis without a remarkable change in mutation spectrum.

The *rnr1Y285F* allele has a modest effect on mutation rate, yet yielded a distinct mutation spectrum especially when paired with *MSH* deletions. In fact, the *rnr1Y285F mshΔ* spectrum closely resembles that of *rnr1Y285A mshΔ*, despite *rnr1Y285A* having a higher skew in dNTP pools, a higher mutation rate (Table 2) and a fitness defect (Fig. 1). We conclude that even modest skewed increases in dNTPs (*rnr1Y285F*) result in distinct error accumulation, likely due to both a decrease in selectivity of the replicative polymerases, and to an increase in efficient mismatch extension at the expense of proofreading (Kumar *et al*. 2011b; Watt *et al*. 2016). Fig. 8A illustrates an example of two different positions in the *rnr1Y285A msh6Δ* background that are predicted to be mutated at increased frequencies via this mechanism.

**Figure 8.**
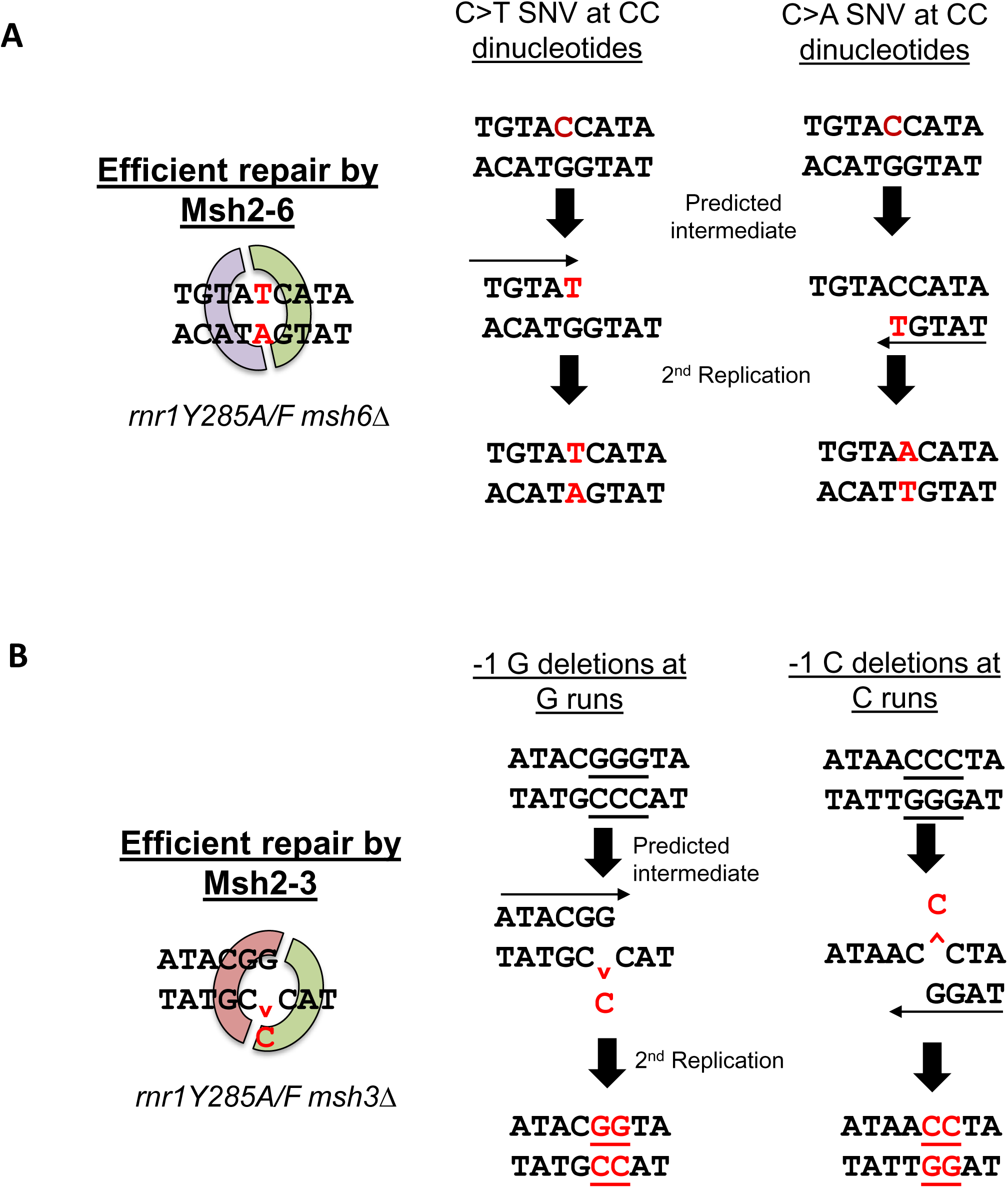
Mechanisms of mutagenesis for the incorporation of errors specific to either Msh2-Msh6 or Msh2-Msh3 repair. The mutated base of interest is represented in red. (A). Two examples of sequence context surrounding CC dinucleotides from Fig. 6B, where mis-insertion is due to the nucleotide in excess in *rnr1Y285F/A* backgrounds. These errors are efficiently repaired by Msh2-Msh6. This specificity becomes apparent when *msh6Δ* is paired with *rnr1Y285F/A* alleles. (B) Two examples of sequence context from Figs. 7A and 7D, where mis-alignment events occur due to the severely limiting amount of dGTP in *rnr1Y285F/A* genetic backgrounds. The run where the deletion occurred is underlined in black. These single base G/C deletions are efficiently repaired by Msh2-Msh3, but not Msh2-Msh6, a previously unidentified specificity of the repair complex.

The increase in G/C deletions in both *rnr1Y285F* and *rnr1Y285A* genotypes can be explained by limiting levels of dGTP (Fig. 8B). dGTP levels are limiting in both yeast and mammalian cells (Chabes *et al*. 2003; Wilson *et al*. 2011; Mathews 2015) and notably led to distinct patterns of mutagenesis when the concentration decreases further, relative to the increase in the other 3 nucleotides, in *rnr1Y285F* and *rnr1Y285A* and even in *rnr1D57N*. This effect is exacerbated by the loss of MMR, especially Msh2- Msh3. Notably, *rnr1* alleles that cause increases in dGTP are also found to be extremely mutagenic (Schmidt *et al*. 2019), highlighting the importance of maintaining the proper level and relative abundance of dGTP.

### New insights into MMR specificity

Msh2-Msh3 and Msh2-Msh6 have separate but overlapping DNA substrate specificities, leading to an expanded repertoire of repair. Previous studies, using a variety of reporter assays, have demonstrated that both can recognize and direct repair of small insertions and deletions (Sia *et al*. 1997; Flores-rozas AND KOLODNER 1998; Harfe AND JINKS-ROBERTSON 1999; Kunkel AND ERIE 2005), but we still have an incomplete understanding of the mechanistic differences in Msh2-Msh3- versus Msh2- Msh6-directed repair. The deep sequencing reported here highlights previously unreported specificities for these pathways and provides new information about sequence context effects on MMR. Previous studies used single-strand oligonucleotide transformation efficiency to define Msh2- Msh3 and Msh2-Msh6 activities (Kow *et al*. 2007; Romanova AND CROUSE 2013). This approach indicated that Msh2-Msh6 preferentially corrects insertions, while Msh2-Msh3 preferentially corrects deletions. We did not observe this bias in our data. Although insertions were relatively infrequent in our data sets, *msh3Δ* and *msh6Δ* exhibited similar levels of insertion events (Fig. 2) although *msh3Δ* exhibited more >1 bp insertions.

By altering dNTP pools, we altered the frequency and types of replication errors generated by DNA polymerases, revealing new substrate specificities for Msh2-Msh3 and Msh2-Msh6. It was striking how the mutation profiles for *msh3Δ* and *msh6Δ* clustered into such distinct groups, in terms of both the types of variants observed and the sequence contexts in which they occurred. While G/C SNVs are common in wild-type backgrounds, G/C-1 deletions are relatively rare, making it difficult to determine the relative efficiencies of MSH complexes in repairing this type of error. However, elevating the dNTP pools to any extent increased the proportion of G/C-1 deletions, allowing us to assess MMR efficacy in their repair. In particular, the meta-analysis of our dataset showed most single base deletions in *rnr1Y285F*, *rnr1Y285A*, and *msh3Δ* occurred in G/C rich contexts, especially homopolymeric runs (Fig. 7, 8B). The double mutants *rnr1Y285F msh3Δ* and *rnr1Y285A msh3Δ* exhibited mutation profiles that were completely dominated by G/C-1 deletions. Therefore Msh2- Msh6 was not able to compensate for the loss of Msh2-Msh3 for repair of G/C-1 deletions within G/C-rich sequence contexts, despite the fact that both complexes have been implicated in directing repair of single base deletions (Meier *et al*. 2018). This suggests a previously unexplored role of Msh2-Msh3 in promoting replication fidelity within G/C-rich genomic regions. It will be interesting to see how different MLH complexes contribute to this specificity. Previous work has indicated that two MLH complexes, Mlh1-Pms1 and Mlh1-Mlh3, are important for repair of deletion mutations (Romanova AND CROUSE 2013).

Similarly, SNVs at GG/CC dinucleotides were more prevalent in *rnr1D57N* and *rnr1Y285F* backgrounds than in wild-type (Fig. 6, 7A). We demonstrated that mutations in these G/C-rich patches were enriched in *msh2Δ* and *msh6Δ* backgrounds, but depleted in *msh3Δ*, indicating that Msh2-Msh6 directs more efficient repair of mutations in GG/CC dinucleotides than Msh2-Msh3. GG dinucleotides are mutated as a signature mutation in colorectal cancer (Rubin AND GREEN 2009), cancers which are defined by defects in MMR. Mutations at CC dinucleotides are also found in human lung cancer (Greenman *et al*. 2007; Lee *et al*. 2010) and could be generated due to defects in MMR and elevations in dNTP levels in combination. Interestingly, the combination of the RNR R2 subunit and deletion of *MSH6* caused a synergistic increase in lung carcinogenesis in a mouse model, although no link with altered dNTP pools was established (Xu *et al*. 2008).

Sequence context analysis indicated that repetitive homoplymeric sequences were the strongest predictor of both in/dels and SNVs. Previous work found specific sequence context effects for human Msh2-Msh6 (hMsh2-hMsh6) activation *in vitro* (Mazurek *et al*. 2009). Mazurek et al (Mazurek *et al*. 2009) found that mispairs surrounded by symmetric 3’ purines were preferentially bound by hMsh2-hMsh6. These substrates also enhanced activation of hMsh2-hMsh6 ATPase activity. Both effects were predictive of enhanced repair and were hypothesized to increase the flexibility of the DNA substrate to allow efficient MSH-DNA complex formation. Our deep sequencing of *can1* did not reveal a strong Msh2-Msh6 bias for these sequence contexts. There was a subset of mispairs surrounded by 3’ purines (trinucleotide contexts e, g, m and o) that were enhanced in the absence of *MSH6*, but the effect was relatively small (∼2-fold increases) and was not systematic. In all genotypes, the greatest predictor of an SNV appeared to be the presence of dinucleotide repeat within the trinucleotide context (GG, CC, TT or AA). As previously noted (Mazurek *et al*. 2009), we observed no distinct broader sequence context implicated in directing MMR beyond homopolymeric runs (Figs. 5-7) although we did note the presence of surrounding A’s in our motif analysis, which may be important for increasing DNA flexibility and bending by MSH complexes.

There are few strand specific effects from altering dNTP levels(Buckland *et al*. 2014), but numerous studies have indicated that MMR is more efficient on the lagging strand (Pavlov *et al*. 2003; Kow *et al*. 2007; Lujan *et al*. 2012). However, Msh2-Msh3 does not appear to have a lagging strand bias and may, in fact, preferentially act on the leading strand (Kow *et al*. 2007). This leads us to hypothesize that Msh2-Msh3 may have greater specificity for lagging strand DNA repair. Our targeted sequencing approach could be applied to strains with *CAN1* in the reverse orientation to explore the differential activity and/or specificity of Msh2-Msh3 and Msh2-Msh6 on leading versus lagging strands. This may be due to distinct interactions of Msh2-Msh3 versus Msh2-Msh6 with MLH complexes (Kow *et al*. 2007; Iyer *et al*. 2010; Kadyrova *et al*. 2020) or PCNA (Lau *et al*. 2002; Iyer *et al*. 2010).

Pairing elevations in dNTP levels with MMR deletions led to increased mutation rates and distinct mutation spectra, similar to previous observations specifically with *msh2Δ*. However, when measuring the rate of canavanine resistance in *rnr1Y285A msh2Δ* and *rnr1Y285A msh6Δ* backgrounds, we observed substantially lower colony numbers under permissive conditions, indicating reduced fitness even under conditions when all cells should be able to grow. Reduced fitness of *rnr1Y285A msh2Δ* and *rnr1Y285A msh6Δ* was also seen in SGA analysis and in tetrad dissections, consistent with the phenomenon of error extinction where the threshold of mutation rate that allows wild type cellular fitness is surpassed (Williams *et al*. 2013; Herr *et al*. 2014; Schmidt *et al*. 2017). A similar growth defect in *rnr1Y285A msh2Δ* was noted in previous work (Watt *et al*. 2016) and other *rnr1* alleles combined with *msh2Δ* also exhibited growth defects (Schmidt *et al*. 2019). We expect that the fitness defects we observe in the absence of selection would also reduce the number of cells that are able to grow under the selective conditions of mutation rate experiments, resulting in mutation rates that are underestimates (Table 2). Our results support a model in which MMR protects cellular fitness in the presence of *rnr1Y285A*. This could be by reducing the level of mutagenesis to avoid mutation-induced extinction (Herr *et al*. 2014). Alternatively, it is possible that the *rnr1Y285A msh2Δ* combination results in a cell cycle defect, as observed with *pol3-01* (Datta *et al*. 2000). In this context, it is intriguing that the SGA screens identified synthetic effects between *rnr1Y285F/A* and several genes involved in mitosis.

### Application to mutation signatures in human cancer

The *msh2Δ, msh3Δ* and *msh6Δ* mutation spectra all had features of MMR deficient human tumor samples (Alexandrov *et al*. 2013). Notably, the *msh6Δ* spectrum in our study closely resembles that of *Msh6^-/-^* in a HAP1 cell line (Zou *et al*. 2018). The percentage of substitutions and IDLs in the *msh6*Δ mutation spectrum is consistent with what is seen in a *Msh6^-/-^* cell line. While T>C variants did not dominate the yeast *msh6Δ* mutation signature as it did in the *Msh6^-/-^* cell line, the overall frequency and proportion of T>C changes did increase significantly. These data indicate that mutation signatures developed through defined mutations and deletions in *S. cerevisiae* will be broadly applicable to mammalian systems. Mutation signatures from *C. elegans* also resemble human cancer signatures, albeit with some minor discrepancies (Meier *et al*. 2014; Meier *et al*. 2018). We note that *C. elegans* lacks a *MSH3* homolog, which is present in both yeast and humans (Denver *et al*. 2005).

Mutation signatures observed in human cancers are routinely used to predict mechanisms of tumorigenesis. We compared our SNV trinucleotide context profiles with the COSMIC single base substitution (SBS) dataset (Alexandrov *et al*. 2020), by hierarchical cluster analysis (Fig. S4, Table S19). Overall, the C>A and C>T changes appeared to drive the clustering. Our samples formed a distinct cluster, which included COSMIC signature SBS32, a mutation profile associated with azathioprine treatment (Fig. S4, box I). The COSMIC cluster most correlated with our samples (Fig. S4, box II, Table S20) included SBS36 (associated with defective base excision repair), SBS38 (unknown association but found only in UV light-associated melanomas), SBS18 (possibly resulting from reactive oxygen species damage), SBS24 (associated with aflatoxin exposure), SBS29 (found in tobacco-chewing related cancers) and SBS52, SBS45 and SBS49 (noted as possible sequencing artefacts).

SBS6 and SBS15 are both most highly correlated with *rnr1Y285F* and *rnr1D57N* paired with *msh2Δ* or *msh6Δ* (Table S20). SBS6 most commonly occurs in colorectal and uterine cancers and is associated with defective MMR. It is possible that elevations in dCTP and dTTP contribute to this mutation signature in human cancers leading to synergistic increases in these types of mutations in the absence of certain MMR genes. SBS15 is characterized by C>T SNVs within NpCpG sequences in lung and stomach cancers with unknown origin. The origin of these cancers could be in part, due to defects in MMR coupled with skewed increases in dCTP and dTTP, consistent with the increased frequency of these SNVs at dinucleotides observed in *rnr1Y285F/A msh6Δ* backgrounds. SBS20 also correlated well with *rnr1Y285F* and *rnr1D57N* combined with *msh2Δ* or *msh6Δ.* This signature is associated with combined MMR and *POLD1* mutations (Table S20). This indicates that reduced replication fidelity resulting from either increased dNTP pools or decreased proofreading combine with MMR defects to generate similar mutation signatures.

In addition, we also compared mutation signatures from our study to the COSMIC insertion and deletion (ID) signatures (Alexandrov *et al*. 2020). A major caveat to comparing *CAN1* ID mutations to human mutation signatures is the lack of homopolymeric sequences within *CAN1*. Nonetheless, within the sequence contexts available, we noted increased G/C single base deletions in G/C context was present in COSMIC ID signatures ID3, ID7, ID9 and ID15. The proposed aetiology of ID Signature 3 is cigarette smoking, ID7 is MMR defects and ID9 and ID15 have unknown etiology. Elevated dNTP levels have not been part of a clinical diagnosis, but skewed increases in dNTPs likely also contribute to these signatures of unknown aetiology. Our results highlight the importance of considering altered dNTP pools and combinatorial effects of genetic backgrounds, when defining the source of tumor mutation signatures.

## Supporting information

Tables S1 and S2_strains and plasmids

Tables S3-S9_SGA data

Table S10_Mutations

Tables S11-S20_ Correlations and hot spots

Table S21_Description of samples

Table S22 _ all variants

## Acknowledgements

We gratefully acknowledge Dr. Anastasia Baryshnikova for her help with the SAFE analysis. N.A.L. is a University at Buffalo Presidential Scholar. This work was supported by the American Cancer Society (RSG-14-2350-01) to J.A.S. J.A.S. is an ACS Research Scholar. J.A.S. is also grateful for support from the University at Buffalo’s Genome, Environment and Microbiome Community of Excellence. This work was also supported by a grant from the Canadian Institutes of Health Research (FDN-159913 to GWB).

