## Supplementary figures for "Complex Mutation Profiles in Mismatch Repair and Ribonucleotide Reductase Mutants Reveal Novel Repair Substrate Specificity of MutS Homolog (MSH) Complexes"

#### Supplementary Figure Legends

**Figure S1. Absolute variant frequency is consistent with selection at *CAN1.* (A)** The absolute variant frequency was calculated by taking the sum of variant frequencies in each genotype and averaging for all biological replicates sequenced in a genotype. Error bars represent the standard deviation between biological replicates within a genotype. The number of total biological replicates sequenced varied by genotype; numbers are displayed in Table S3. Samples grown under permissive conditions, from four genotypes, are graphed in peach. All other samples grown under selection are graphed in turquoise. **(B)** The average absolute variant frequency before (red) and after (turquoise) the permissive variant filter was applied.

**Figure S2. Hierarchical cluster analysis using Spearman rank correlation on all individual samples in our study.** Boxed in black are notable clusters where biological replicates from the same genotypes cluster together. Throughout, biological replicates tended to cluster. Similarly, *rnr1Y285F pGAL-RNR1* and *rnr1Y285A pGAL-RNR1* strains clustered with *rnr1Y285F* and *rnr1Y285A* strain, respectively.

**Figure S3. Mutation spectra comparison between rnr1 alleles and *rnr1-pGAL-RNR1* counterparts.**

**(A)** The SNV spectra normalized out of total SNVs and **(B)** total variants. **(C)** The deletion spectra normalized out of total deletions and **(D)** total variants. **(E)** The insertion spectra normalized out of total insertions and **(F)** total variants.

**Figure S4. Hierarchical cluster analysis of mutation signatures from this study compared with COSMIC SBS signatures.** The single base substitutions (SBS) COSMIC signatures from GRCh38 (v3.2- March 2021, https://cancer.sanger.ac.uk/signatures/downloads/) were combined with the normalized SNVs in trinucleotide context (**Fig. 4).** Our samples formed a distinct cluster, which included COSMIC signature SBS32, a mutation profile associated with azathioprine treatment (**box I**). A second cluster (**box II**) correlated well with several of our samples (**Table S20**).

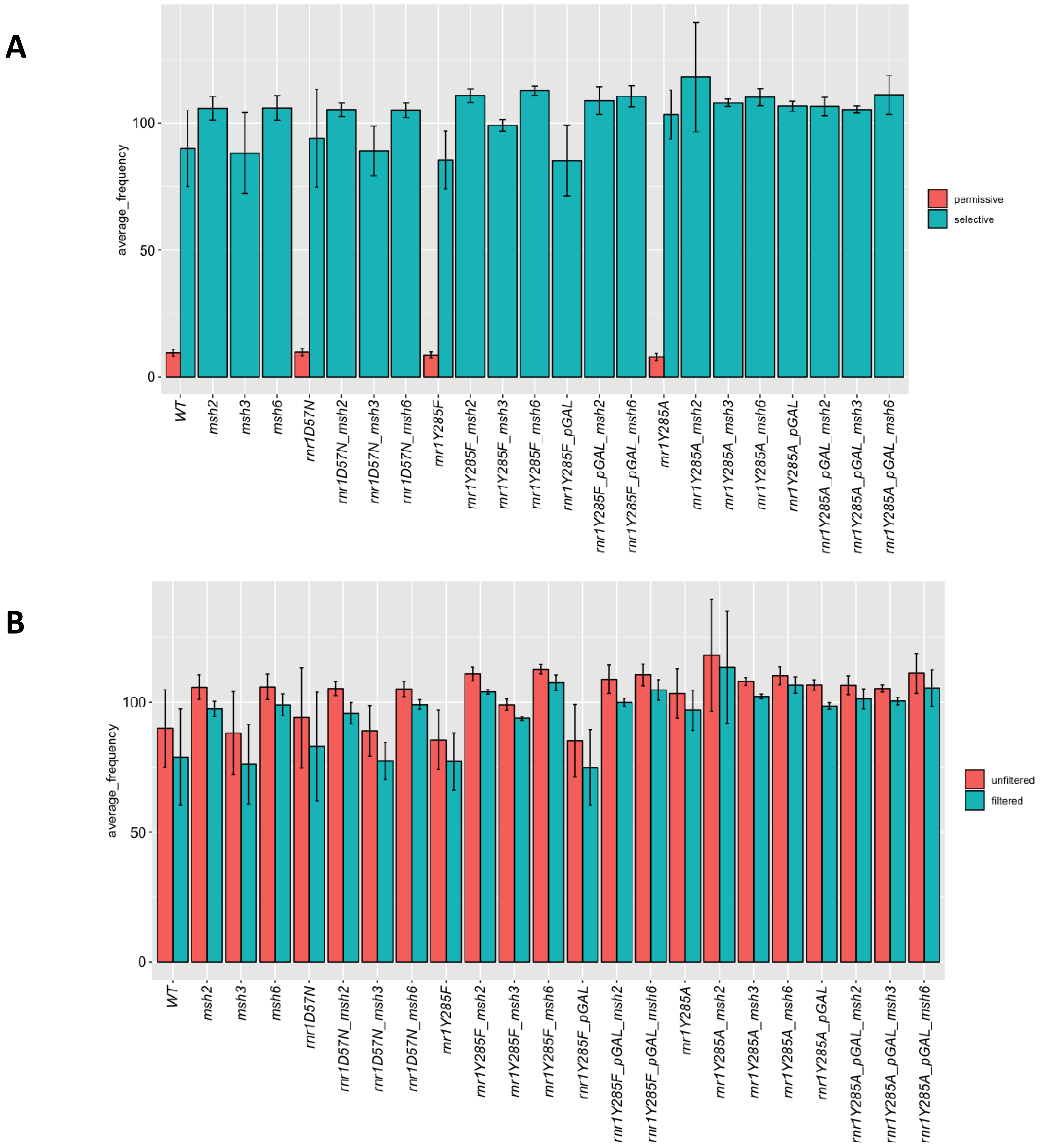
A
average_frequency
0
50
100
permissive
selective
WT
msh2
msh3
msh6
rnr1D57N
rnr1D57N_msh2
rnr1D57N_msh3
rnr1D57N_msh6
rnr1Y285F
rnr1Y285F_msh2
rnr1Y285F_msh3
rnr1Y285F_msh6
rnr1Y285F_pGAL
rnr1Y285F_pGAL_msh2
rnr1Y285F_pGAL_msh6
rnr1Y285A
rnr1Y285A_msh2
rnr1Y285A_msh3
rnr1Y285A_msh6
rnr1Y285A_pGAL
rnr1Y285A_pGAL_msh2
rnr1Y285A_pGAL_msh3
rnr1Y285A_pGAL_msh6
B
average_frequency
0
50
100
unfiltered
filtered
WT
msh2
msh3
msh6
rnr1D57N
rnr1D57N_msh2
rnr1D57N_msh3
rnr1D57N_msh6
rnr1Y285F
rnr1Y285F_msh2
rnr1Y285F_msh3
rnr1Y285F_msh6
rnr1Y285F_pGAL
rnr1Y285F_pGAL_msh2
rnr1Y285F_pGAL_msh6
rnr1Y285A
rnr1Y285A_msh2
rnr1Y285A_msh3
rnr1Y285A_msh6
rnr1Y285A_pGAL
rnr1Y285A_pGAL_msh2
rnr1Y285A_pGAL_msh3
rnr1Y285A_pGAL_msh6

**Figure S1.**

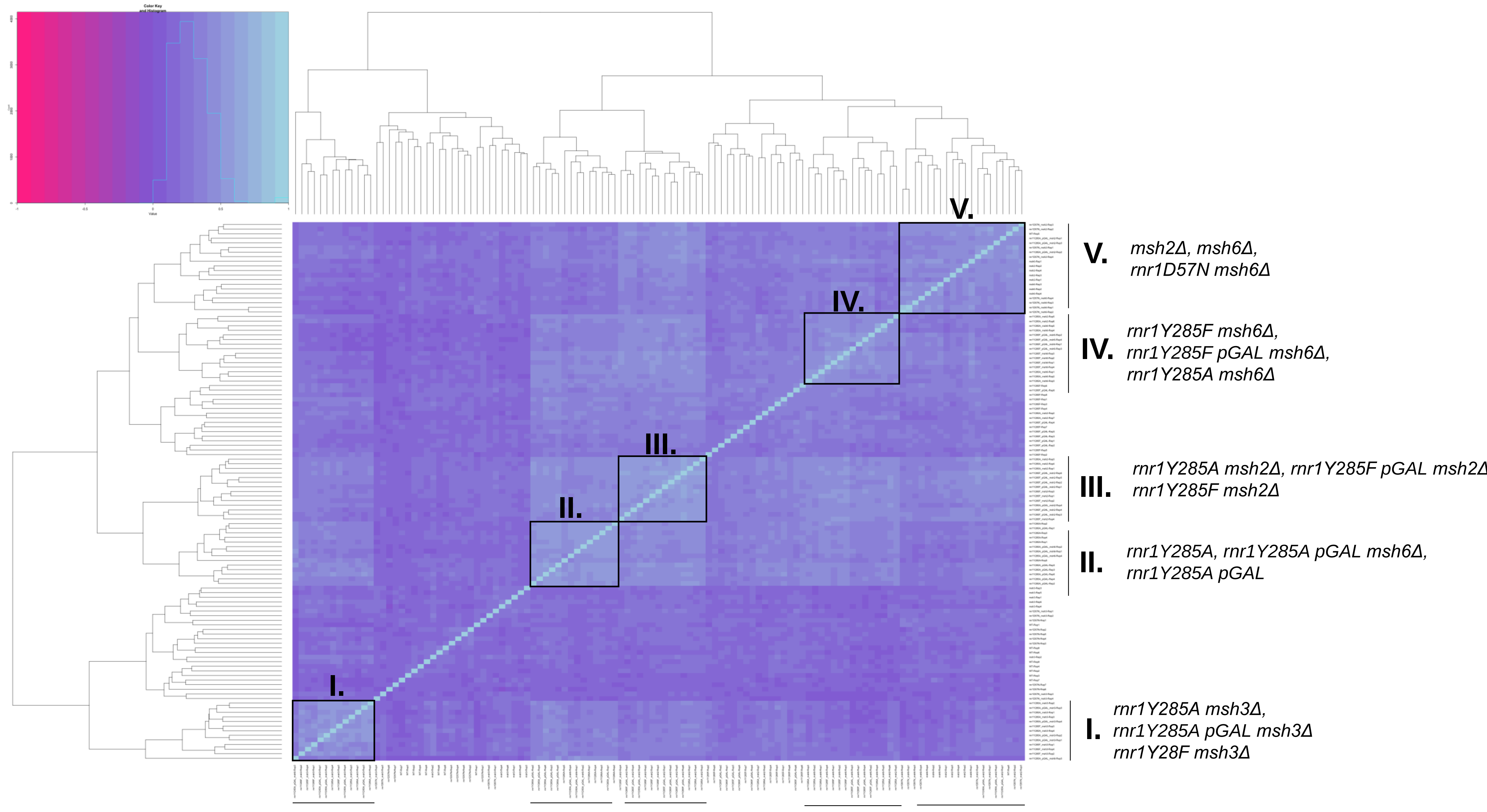

V.
IV.
III.
II.
I.
V. msh2Δ, msh6Δ,
rnr1D57N msh6Δ
IV. rnr1Y285F msh6Δ,
rnr1Y285F pGAL msh6Δ,
rnr1Y285A msh6Δ
III. rnr1Y285A msh2Δ, rnr1Y285F pGAL msh2Δ
rnr1Y285F msh2Δ
II. rnr1Y285A, rnr1Y285A pGAL msh6Δ,
rnr1Y285A pGAL
I. rnr1Y285A msh3Δ,
rnr1Y285A pGAL msh3Δ
rnr1Y28F msh3Δ


**Figure S2**

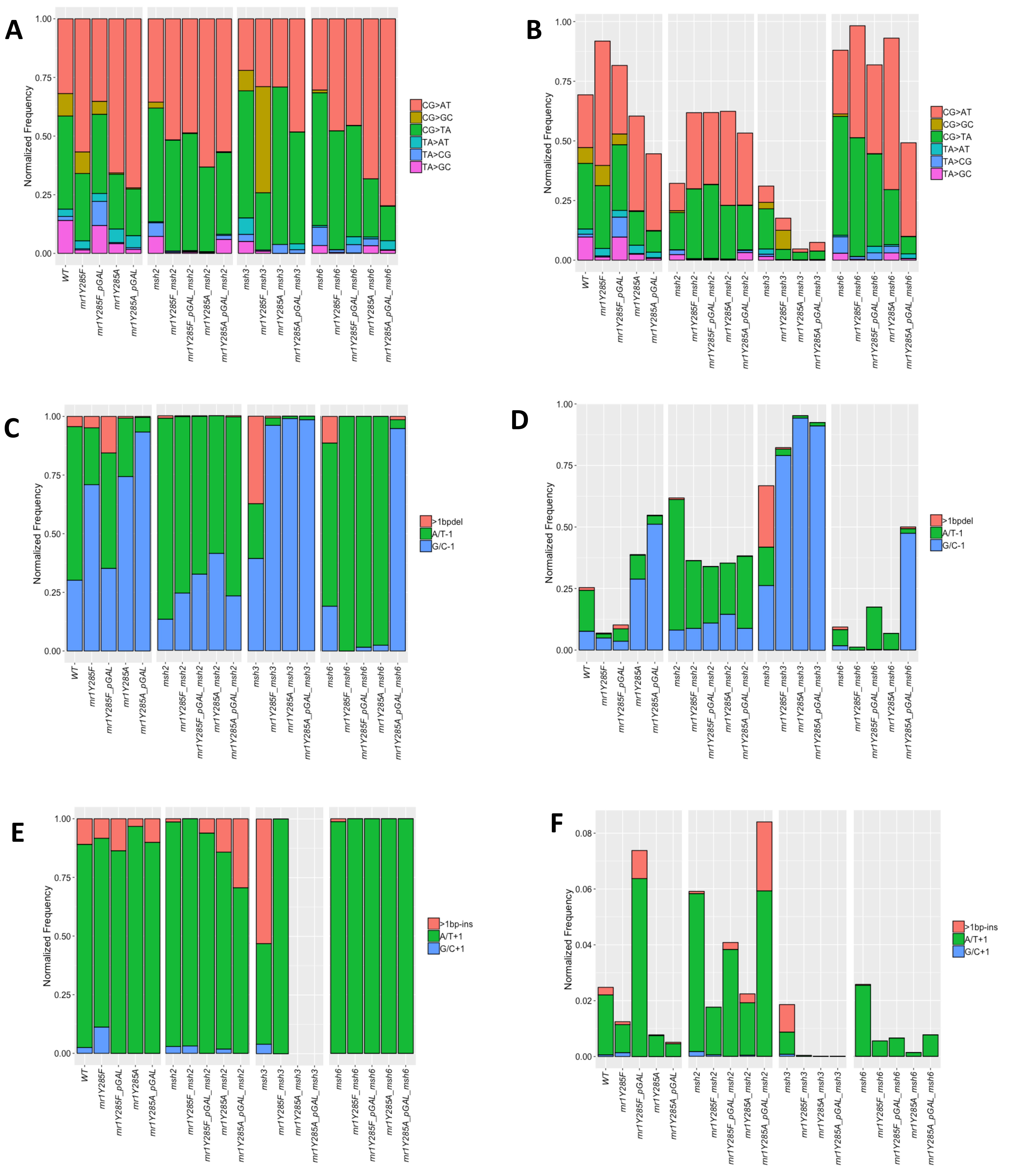
A
Normalized Frequency
1.00
0.75
0.50
0.25
0.00
CG>AT
CG>GC
CG>TA
TA>AT
TA>CG
TA>GC
WT
mr1Y285F
mr1Y285F_pGAL
mr1Y285A
mr1Y285A_pGAL
msh2
mr1Y285F_msh2
mr1Y285F_pGAL_msh2
mr1Y285A_msh2
mr1Y285A_pGAL_msh2
msh3
mr1Y285F_msh3
mr1Y285A_msh3
mr1Y285A_pGAL_msh3
msh6
mr1Y285F_msh6
mr1Y285F_pGAL_msh6
mr1Y285A_msh6
mr1Y285A_pGAL_msh6
B
C
>1bpdel
A/T-1
G/C-1
D
E
>1bp-ins
A/T+1
G/C+1
F
0.08
0.06
0.04
0.02

Figure S3

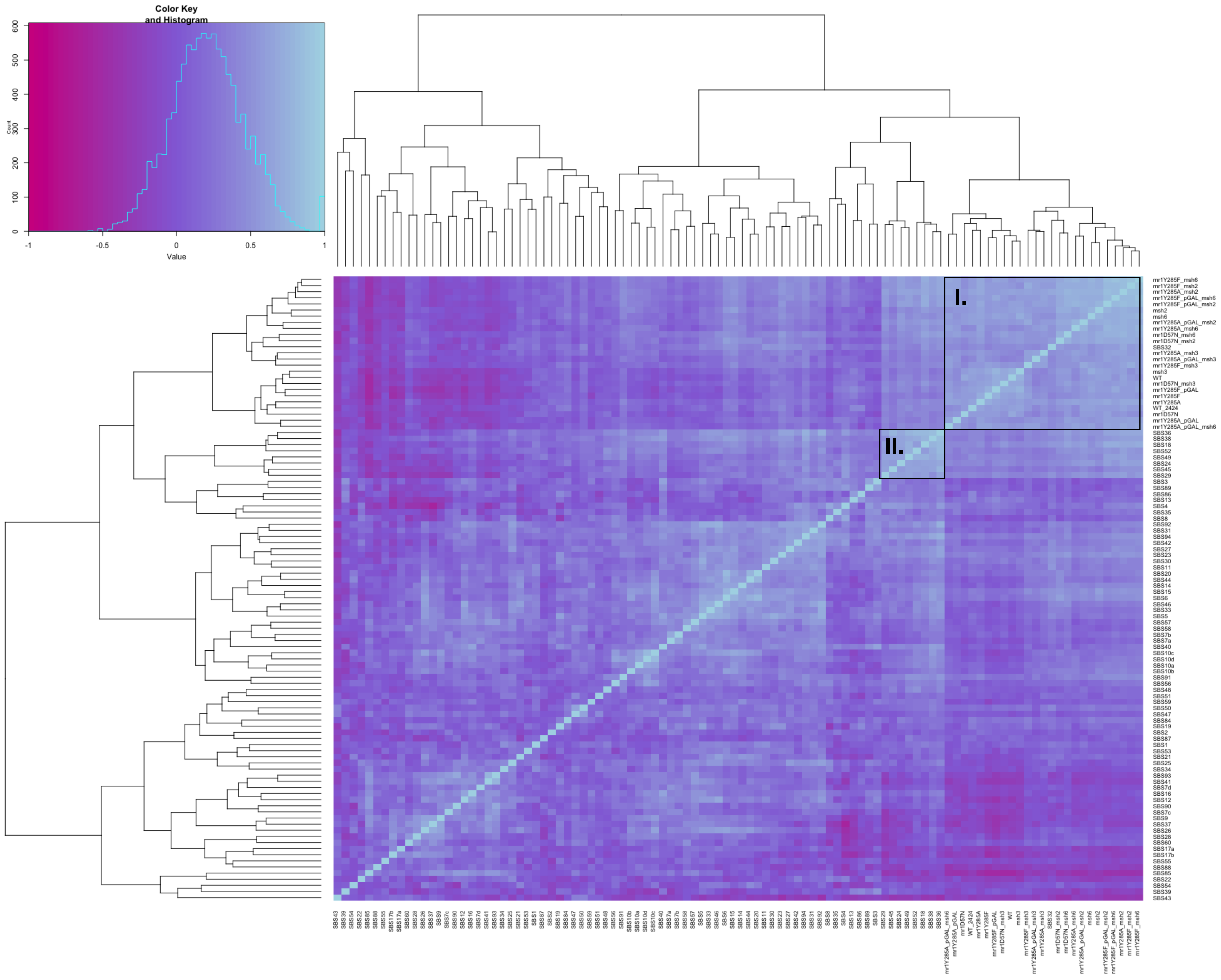


### Figure S4
