## Supplementary material for "Complex Mutation Profiles in Mismatch Repair and Ribonucleotide Reductase Mutants Reveal Novel Repair Substrate Specificity of MutS Homolog (MSH) Complexes": Tables S1 and S2_strains and plasmids

**Supplementary Table S1. Strains used in this study**

Except as indicated, all strains are in the W303 background:

*CAN1 ade2-1 his3-11,15 leu2-3,112 trp1-1 ura3-1 RAD5+* or *ade2-11 his3-11,15 leu2-3,112 trp1-1 URA3 RAD5+*

| JSY# | Other name | Relevant Genotype | Phenotype | Source |
|---|---|---|---|---|
| JSY13 | EAY1995/4C | *MAT***a** *RNR1 CAN1 ade2-1 his3-11,15 leu2-3,112 trp1-1 ura3-1 RAD5+* | Wildtype | Chabes et al., 2003<br>Xu et al., 2008 |
| JSY2424 | AC402 | *RNR1 ade2-11 his3-11,15 leu 2-3112 trp1-1 URA3 RAD5+* | Wildtype | Chabes et al., 2003 |
| JSY22 | EAY2004 | *msh2Δ* | loss of MMR | Xu et al., 2008 |
| JSY23 | EAY2005 | *msh2Δ* | loss of MMR | Xu et al., 2008 |
| JSY15 | EAY1997 | *msh3Δ* | loss of Msh2-Msh3 directed repair | Xu et al., 2008 |
| JSY16 | EAY1998 | *msh3Δ* | loss of Msh2-Msh3 directed repair | Xu et al., 2008 |
| JSY28 | EAY2010 | *msh6Δ* | loss of Msh2-Msh6 directed repair | Xu et al., 2008 |
| JSY29 | EAY2011 | *msh6Δ* | loss of Msh2-Msh6 directed repair | Xu et al., 2008 |
| JSY14 | EAY1995/8C | *rnr1D57N* | 2-fold balanced elevation in dNTPs | Chabes et al., 2003 |
| JSY25 | EAY2007 | *rnr1D57N msh2Δ* | Predicted 2-fold balanced elevation in dNTPs, loss of Msh2 directed repair | Xu et al., 2008 |
| JSY26 | EAY2008 | *rnr1D57N msh2Δ* | Predicted 2-fold balanced elevation in | Xu et al., 2008 |

| | | | | |
|---|---|---|---|---|
| | | | dNTPs, loss of Msh2 directed repair | |
| JSY20 | EAY2002 | *rnr1D57N msh3Δ* | Predicted 2-fold balanced elevation in dNTPs, loss of Msh3 directed repair | Xu et al., 2008 |
| JSY21 | EAY2003 | *rnr1D57N msh3Δ* | Predicted 2-fold balanced elevation in dNTPs, loss of Msh3 directed repair | Xu et al., 2008 |
| JSY32 | EAY2014 | *rnr1D57N msh6Δ* | Predicted 2-fold balanced elevation in dNTPs, loss of Msh6 directed repair | Xu et al., 2008 |
| JSY33 | EAY2015 | *rnr1D57N msh6Δ* | Predicted 2-fold balanced elevation in dNTPs, loss of Msh6 directed repair | Xu et al., 2008 |
| JSY2420, JSY2421 | DK3A, DK3C | *rnr1Y285F URA3-pGAL-RNR1* | 2-fold increase in dCTP and dTTP | Kumar et al., 2010 |
| JSY2422, JSY2423 | DK8A, DK8E | *rnr1Y285A URA3-pGAL-RNR1* | 20-fold increase in dCTP and dTTP | Kumar et al., 2010 |
| JSY3811,JSY3865,JSY3866 | JSY3811,JSY3865,JSY3866 | *rnr1Y285F* | Predicted 2-fold increase in dCTP and dTTP | Lamb et al, 2021 |
| JSY3818-3820 | JSY3818<br>JSY3819 | *rnr1Y285F msh6Δ* | Predicted 2-fold increase in dCTP and dTTP, loss of Msh6 directed repair | This Study |
| JSY3803-3805 | JSY3803 | *rnr1Y285F msh6Δ URA3-pGAL-RNR1* | Predicted 2-fold increase in dCTP and | This Study |

| | | | | |
|---|---|---|---|---|
| | | | dTTP, loss of Msh6 directed repair | |
| JSY3812-3814 | JSY3812<br>JSY3813 | *rnr1Y285F msh2Δ* | Predicted 2-fold increase in dCTP and dTTP, loss of Msh2 directed repair | This Study |
| JSY3223-3225 | JSY3223<br>JSY3225 | *rnr1Y285F msh2Δ URA3-pGAL-RNR1* | Predicted 2-fold increase in dCTP and dTTP, loss of Msh2 directed repair | This Study |
| JSY3815-3817 | JSY3815<br>JSY3816 | *rnr1Y285F msh3Δ* | Predicted 2-fold increase in dCTP and dTTP, loss of Msh3 directed repair | This Study |
| JSY3868-<br>3869 | JSY3868<br>JSY3869 | *rnr1Y285A* | Predicted 2-fold increase in dCTP and dTTP | Lamb et al, 2021 |
| JSY3882 | JSY3882 | *rnr1Y285A msh6Δ* | Predicted 20-fold increase in dCTP and dTTP, loss of Msh6 directed repair | This Study |
| JSY3809-3810 | JSY3809<br>JSY3810 | *rnr1Y285A msh6Δ URA3-pGAL-RNR1* | Predicted 20-fold increase in dCTP and dTTP, loss of Msh6 directed repair | This Study |
| JSY3872-3873 | JSY3872<br>JSY3873 | *rnr1Y285A msh2Δ* | Predicted 20-fold increase in dCTP and dTTP, loss of Msh2 directed repair | This Study |
| JSY3228-3229 | JSY3228<br>JSY3229 | *rnr1Y285A msh2Δ URA3-pGAL-RNR1* | Predicted 20-fold increase in dCTP and dTTP, loss of Msh2 directed | This Study |

| | | | | |
|---|---|---|---|---|
| | | | repair | |
| JSY3876-3879 | JSY3876<br>JSY3879 | *rnr1Y285A msh3Δ* | Predicted 20-fold increase in dCTP and dTTP, loss of Msh3 directed repair | This Study |
| JSY3650-3682 | JSY3650<br>JSY3682 | *rnr1Y285A msh3Δ URA3-pGAL-RNR1* | Predicted 20-fold increase in dCTP and dTTP, loss of Msh3 directed repair | This Study |
| JSY1379<br>S288C (BY) | | *MATα rnr1-D57N::natMX can1Δ::STE2pr-sp_his5+ leu2Δ0 his3Δ1 ura3Δ0 met15Δ0 lyp1Δ LYS2* | Predicted 2-fold balanced elevation in dNTPs | This study |
| JSY3706<br>S288C (BY) | | *MATα rnr1-Y285F::natMX can1Δ::STE2pr-sp_his5+ leu2Δ0 his3Δ1 ura3Δ0 met15Δ0 lyp1Δ LYS2* | Predicted 2-fold increase in dCTP and dTTP | This study |
| JSY3709<br>S288C (BY) | | *MATα rnr1Y285A::natMX can1Δ::STE2pr-Sp_his5 leu2Δ0 his3Δ1 ura3Δ0 met15Δ0 lyp1Δ LYS2* | Predicted 20-fold increase in dCTP and dTTP | This study |
| yGWB2487<br>S288C (BY) | | *MAT***a** *msh2Δ0::kanMX leu2Δ0 his3Δ1 ura3Δ0 met15Δ0* | | This study |
| yGWB7124<br>S288C (BY) | | *MAT***a** *msh3Δ0::kanMX leu2Δ0 his3Δ1 ura3Δ0 met15Δ0* | | This study |
| yGWB7125<br>S288C (BY) | | *MAT***a** *msh6Δ0::kanMX leu2Δ0 his3Δ1 ura3Δ0 met15Δ0* | | This study |

**Supplementary Table S2. Primers used in this study**

| SO# | Name | Sequence- 5'-3' | Description |
|---|---|---|---|
| 750 | MSH2 PRIMER A | CGTATAAACAAAGCCAAAGACAAGT | Amplifying MSH2::Kan MX |
| 751 | MSH2 PRIMER D | ACATCTCTTGTTTATCCCATCCATA | Amplifying MSH2::Kan MX |
| 752 | MSH3 PRIMER A | CCTGTTTTTCCTTTGATGTTTCTAA | Amplifying MSH3::Kan MX |
| 753 | MSH3 PRIMER D | TGATCCATTCCATGATTTTAATTCT | Amplifying MSH3::Kan MX |
| 754 | MSH6 PRIMER A | GTCTCCATTTCCAACTAATGGTATG | Amplifying MSH6::Kan MX |
| 755 | MSH6 PRIMER D | AGCTGAATCATAGGTCAAGAAAATG | Amplifying MSH6::Kan MX |
| 807 | CAN1 reg1 Forward_anchored | TCGTCGGCAGCGTCAGATGTGTATAAGAGACAGCTCCTGTAAAAACAAAAAAAAAAAAAGCG | Sequencing primers with nextera adapters: CAN1-region1 |
| 714 | CAN1 reg1 Reverse | GTCTCGTGGGCTCGGAGATGTGTATAAGAGACAGTAGTACCACCAAGGGCAATC | Sequencing primers with nextera adapters: CAN1-region1 |
| 715 | CAN1 reg2 Forward | TCGTCGGCAGCGTCAGATGTGTATAAGAGACAGAAGCAAAGACATATTGGTAT | Sequencing primers with nextera adapters: CAN1-region2 |
| 716 | CAN1 reg2 Reverse | GTCTCGTGGGCTCGGAGATGTGTATAAGAGACAGATCCATGCGCGCAGTGGAAC | Sequencing primers with nextera adapters: CAN1- |

| | | | |
|---|---|---|---|
| | | | region2 |
| 717 | CAN1 reg3 Forward | TCGTCGGCAGCGTCAGATGTGTATAAGAGACAGTTCAATTTTGGACGTACAAA | Sequencing primers with nextera adapters: CAN1-region3 |
| 718 | CAN1 reg3 Reverse | GTCTCGTGGGCTCGGAGATGTGTATAAGAGACAGACCAGCAGTGATACCAACTA | Sequencing primers with nextera adapters: CAN1-region3 |
| 719 | CAN1 reg4 Forward | TCGTCGGCAGCGTCAGATGTGTATAAGAGACAGCACATTTCAAGGTACTGAAC | Sequencing primers with nextera adapters: CAN1-region4 |
| 720 | CAN1 reg4 Reverse | GTCTCGTGGGCTCGGAGATGTGTATAAGAGACAGTAGGAGCCAACTTGTTCTTT | Sequencing primers with nextera adapters: CAN1-region4 |
| 721 | CAN1 reg5 Forward | TCGTCGGCAGCGTCAGATGTGTATAAGAGACAGCGTATTTTATTTGGTCTATC | Sequencing primers with nextera adapters: CAN1-region5 |
| 722 | CAN1 reg5 Reverse | GTCTCGTGGGCTCGGAGATGTGTATAAGAGACAGAAACCTTGAATAATGATAAT | Sequencing primers with nextera adapters: CAN1-region5 |
| 723 | CAN1 reg6 Forward | TCGTCGGCAGCGTCAGATGTGTATAAGAGACAGCGGCCACATTTATGACGATC | Sequencing primers with nextera adapters: |

| | | | |
|---|---|---|---|
| | | | CAN1-region6 |
| 724 | CAN1 reg6 Reverse | GTCTCGTGGGCTCGGAGATGTGTATAAGAGACAGCTTTCTTTTCGGTGTATGAC | Sequencing primers with nextera adapters: CAN1-region6 |
