## Supplementary material for "Complex Mutation Profiles in Mismatch Repair and Ribonucleotide Reductase Mutants Reveal Novel Repair Substrate Specificity of MutS Homolog (MSH) Complexes": Table S10_Mutations

**Supplementary Table S10. Mutation spectra**

| | WT (9)[a] | | *msh2Δ* (4) | | *msh3Δ* (6) | | *msh6Δ* (4) | |
|---|---|---|---|---|---|---|---|---|
| **Variants** | **Counts[b]** | **Freq.[c]** | **Counts[b]** | **Freq.[c]** | **Counts[b]** | **Freq.[c]** | **Counts[b]** | **Freq.[c]** |
| **CG>AT** | 18.11 | 17.42 | 19.00 | 11.13 | 15.83 | 5.20 | 45.50 | 26.44 |
| **CG>GC** | 7.44 | 5.23 | 1.00 | 0.79 | 8.00 | 2.06 | 4.00 | 1.03 |
| **CG>TA** | 31.89 | 21.68 | 36.75 | 15.18 | 28.50 | 12.81 | 66.50 | 49.32 |
| **TA>AT** | 4.67 | 1.72 | 0.75 | -.13 | 4.50 | 1.66 | 3.00 | 0.66 |
| **TA>CG** | 3.67 | 0.92 | 6.75 | 1.83 | 4.00 | -.71 | 16.00 | 6.79 |
| **TA>GC** | 4.67 | 7.67 | 3.50 | 2.26 | 4.17 | 1.20 | 6.75 | 2.91 |
| **SNV total[d]** | **70.45** | **54.64** | **67.75** | **31.32** | **65.00** | **23.65** | **141.75** | **87.15** |
| **% SNV[e]** | **79.96** | **69.3** | **68.26** | **32.15** | **61.03** | **31.07** | **88.59** | **88.01** |
| **A/T-1** | 8.44 | 13.11 | 18.75 | 51.69 | 10.33 | 11.89 | 9.00 | 6.45 |
| **G/G-1** | 2.56 | 6.04 | 5.00 | 8.00 | 12.17 | 20.03 | 3.00 | 1.77 |
| **>1 bp del** | 2.11 | 0.87 | 1.000 | 0.62 | 12.83 | 18.99 | 1.25 | 1.05 |
| **Deletion total[f]** | **13.11** | **20.02** | **24.75** | **60.30** | **36.33** | **50.91** | **13.25** | **9.28** |
| **% Deletion[g]** | **14.88** | **25.39** | **24.94** | **61.91** | **33.18** | **66.86** | **8.28** | **9.38** |
| **A/T+1** | 2.56 | 1.69 | 5.25 | 5.51 | 2.50 | -.61 | 4.50 | 2.52 |
| **G/G+1** | 0.22 | 0.05 | 0.75 | 0.16 | 0.17 | 0.06 | 0 | 0 |
| **>1 bp insert** | 0.89 | 0.21 | 0.50 | 0.08 | 2.67 | 0.75 | 0.25 | 0.03 |
| **Insertion total[h]** | **3.67** | **1.95** | **6.50** | **5.75** | **5.33** | **1.41** | **4.75** | **2.55** |
| **% Insertion[i]** | **4.17** | **2.47** | **6.55** | **5.91** | **5.01** | **1.85** | **2.97** | **2.58** |
| **MNV** | 0.44 | 1.92 | 0.25 | 0.03 | 0.33 | 0.10 | 0.25 | 0.04 |
| **Replacement** | 0.44 | 0.31 | 0 | 0 | 0.50 | 0.07 | 0 | 0 |
| **Complex total[j]** | **0.88** | **2.23** | **0.25** | **0.03** | **0.83** | **0.17** | **0.25** | **0.04** |
| **% Complex[k]** | **1** | **2.83** | **0.25** | **0.03** | **0.78** | **0.22** | **0.16** | **0.04** |
| **Sum total[l]** | **88.11** | **78.84** | **99.25** | **97.41** | **106.50** | **76.14** | **160** | **99.02** |
| **Counts/Freq** | **1.12** | | **1.02** | | **1.40** | | **1.62** | |

[a] Number of replicates

[b] Sum of unique variants observed within a genotype divided by the number of replicates

[c] Sum of variant frequencies observed within a genotype divided by the number of replicates

[d] Average total number of SNVs per replicate for each genotype

[e] Percentage of total variants that were SNVs

[f] Average total number of deletions per replicate for each genotype

[g] Percentage of total variants that were deletions

[h] Average total number of insertions per replicate for each genotype

[i] Percentage of total variants that were insertions

[j] Average total number of complex mutations per replicate for each genotype

[k] Percentage of total variants that were complex mutations

[l] Sum of each type of mutation

| **Table S10 (cont'd). Mutation spectra** | | | | | | | | |
|---|---|---|---|---|---|---|---|---|
| | ***rnr1D57N* (7)[a]** | | ***rnr1D57N msh2Δ* (4)** | | ***rnr1D57N msh3Δ* (4)** | | ***rnr1D57N msh6Δ* (4)** | |
| **Variants** | **Counts[b]** | **Freq.[c]** | **Counts[b]** | **Freq.[c]** | **Counts[b]** | **Freq.[c]** | **Counts[b]** | **Freq.[c]** |
| **CG>AT** | 14.00 | 13.48 | 12.00 | 4.58 | 14.25 | 7.77 | 31.00 | 25.84 |
| **CG>GC** | 4.71 | 1.40 | 0.25 | 0.05 | 6.25 | 1.72 | 1.75 | 1.36 |
| **CG>TA** | 25.43 | 9.55 | 41.5 | 20.23 | 20.75 | 18.49 | 59.25 | 52.16 |
| **TA>AT** | 2.57 | 1.19 | 0.50 | 0.30 | 3.00 | 6.35 | 3.75 | 0.98 |
| **TA>CG** | 3.00 | 1.03 | 4.50 | 1.66 | 3.25 | 1.38 | 9.00 | 2.27 |
| **TA>GC** | 3.14 | 4.09 | 1.00 | 0.32 | 3.25 | 1.16 | 4.75 | 2.09 |
| **SNV total[d]** | **52.86** | **30.74** | **59.75** | **27.14** | **50.75** | **36.87** | **109.50** | **85.79** |
| **% SNV[e]** | **75.67** | **37.04** | **70.71** | **28.33** | **60.06** | **47.71** | **86.39** | **86.56** |
| **A/T-1** | 6.57 | 12.99 | 13.50 | 49.23 | 11.00 | 10.17 | 8.25 | 3.71 |
| **G/G-1** | 3.14 | 35.34 | 4.50 | 2.81 | 11.75 | 11.87 | 2.50 | 0.56 |
| **>1 bp del** | 2.14 | 1.30 | 0 | 0 | 5.25 | 15.11 | 0.25 | 4.08 |
| **Deletion total[f]** | **11.86** | **49.62** | **18.00** | **52.04** | **28.00** | **37.15** | **11.00** | **8.35** |
| **% Deletion[g]** | **16.97** | **59.80** | **21.30** | **54.32** | **33.14** | **48.08** | **8.68** | **8.43** |
| **A/T+1** | 2.86 | 1.65 | 5.25 | 16.19 | 3.25 | 2.54 | 5.50 | 4.85 |
| **G/G+1** | 0.57 | 0.37 | 0.25 | 0.03 | 1.75 | 0.15 | 0.25 | 0.04 |
| **>1 bp insert** | 0.57 | 0.32 | 1.25 | 0.39 | 0.50 | 0.53 | 0 | 0 |
| **Insertion total[h]** | **4.00** | **2.34** | **6.75** | **16.61** | **5.50** | **3.22** | **5.75** | **4.89** |
| **% Insertion[i]** | **5.73** | **2.82** | **7.99** | **17.34** | **6.51** | **4.17** | **4.54** | **4.93** |
| **MNV** | 0.57 | 0.14 | 0 | 0 | 0.25 | 0.04 | 0.50 | 0.85 |
| **Replacement** | 0.57 | 0.15 | 0 | 0 | 0 | 0 | 0 | 0 |
| **Complex total[j]** | **1.14** | **0.28** | **0** | **0** | **0.25** | **0.04** | **0.50** | **0.09** |
| **% Complex[k]** | **1.64** | **0.35** | **0** | **0** | **0.30** | **0.05** | **0.40** | **0.09** |
| **Sum total[l]** | **69.86** | **82.99** | **84.50** | **95.79** | **84.50** | **77.28** | **126.75** | **99.12** |
| **Counts/Freq.** | **0.84** | | **0.88** | | **1.09** | | **1.28** | |

| Table S10 (cont'd). Mutation spectra | | | | | | | | |
|---|---|---|---|---|---|---|---|---|
| | *rnr1Y285F* (8)[a] | | *rnr1Y285F msh2Δ* (4) | | *rnr1Y285F msh3Δ* (4) | | *rnr1Y285F msh6Δ* (4) | |
| **Variants** | **Counts[b]** | **Freq.[c]** | **Counts[b]** | **Freq.[c]** | **Counts[b]** | **Freq.[c]** | **Counts[b]** | **Freq.[c]** |
| **CG>AT** | 20.25 | 40.21 | 39.75 | 33.14 | 47.25 | 50.42 | 6.00 | 4.78 |
| **CG>GC** | 5.00 | 6.52 | 0.25 | 0.10 | 0 | 0 | 1.50 | 7.48 |
| **CG>TA** | 39.00 | 20.33 | 45.00 | 30.34 | 48.00 | 53.54 | 22.00 | 4.03 |
| **TA>AT** | 3.00 | 2.42 | 0.25 | 0.04 | 0.50 | 0.11 | 0 | 0 |
| **TA>CG** | 1.50 | 0.34 | 1.25 | 0.35 | 2.50 | 1.13 | 0.50 | 0.08 |
| **TA>GC** | 3.00 | 1.04 | 0.75 | 0.26 | 0.50 | 0.44 | 0.50 | 0.16 |
| **SNV total[d]** | **71.75** | **70.86** | **87.25** | **64.23** | **98.75** | **105.64** | **30.50** | **16.53** |
| **% SNV[e]** | **85.67** | **91.83** | **77.56** | **61.75** | **94.95** | **98.26** | **57.28** | **17.61** |
| **A/T-1** | 4.63 | 1.30 | 13.00 | 28.56 | 3.75 | 1.28 | 4.50 | 2.45 |
| **G/G-1** | 4.38 | 3.79 | 7.75 | 9.26 | 0 | 0 | 15.00 | 74.24 |
| **>1 bp del** | 1.25 | 0.26 | 0.50 | 0.13 | 0 | 0 | 3.00 | 0.57 |
| **Deletion total[f]** | **10.25** | **5.35** | **21.25** | **37.95** | **3.75** | **1.28** | **22.50** | **77.27** |
| **% Deletion[g]** | **12.24** | **6.93** | **18.89** | **36.49** | **3.61** | **1.19** | **42.25** | **82.36** |
| **A/T+1** | 1.50 | 0.78 | 3.75 | 1.77 | 1.50 | 0.59 | 0.25 | 0.03 |
| **G/G+1** | 0.13 | 0.11 | 0.25 | 0.06 | 0 | 0 | 0 | 0 |
| **>1 bp insert** | 0.13 | 0.08 | 0 | 0 | 0 | 0 | 0 | 0 |
| **Insertion total[h]** | **1.75** | **0.96** | **4.00** | **1.83** | **1.50** | **0.59** | **0.25** | **0.03** |
| **% Insertion[i]** | **2.09** | **1.25** | **3.56** | **1.76** | **1.44** | **0.55** | **0.47** | **0.03** |
| **MNV** | 0 | 0 | 0 | 0 | 0 | 0 | 0 | 0 |
| **Replacement** | 0 | 0 | 0 | 0 | 0 | 0 | 0 | 0 |
| **Complex total[j]** | **0** | **0** | **0** | **0** | **0** | **0** | **0** | **0** |
| **% Complex[k]** | **0** | **0** | **0** | **0** | **0** | **0** | **0** | **0** |
| **Sum total[l]** | **83.75** | **77.16** | **112.50** | **104.01** | **104.00** | **107.51** | **53.25** | **93.82** |
| **Counts/Freq** | **1.09** | | **1.08** | | **0.97** | | **0.57** | |

| Table S10 (cont'd). Mutation spectra | | | | | | |
|---|---|---|---|---|---|---|
| | *rnr1Y285F pGAL-RNR1* (6)[a] | | *rnr1Y285F pGAL-RNR1 msh2Δ* (6) | | *rnr1Y285F pGAL-RNR1 msh6Δ* (4) | |
| **Variants** | **Counts[b]** | **Freq.[c]** | **Counts[b]** | **Freq.[c]** | **Counts[b]** | **Freq.[c]** |
| **CG>AT** | 35.00 | 21.51 | 34.83 | 30.05 | 41.00 | 38.98 |
| **CG>GC** | 11.33 | 3.38 | 0.17 | 0.15 | 0.50 | 0.13 |
| **CG>TA** | 47.50 | 20.64 | 44.67 | 30.85 | 34,50 | 40.57 |
| **TA>AT** | 7.33 | 2.08 | 1.17 | 0.22 | 1.50 | 2.88 |
| **TA>CG** | 2.50 | 6.26 | 1.00 | 0.19 | 3.25 | 3.06 |
| **TA>GC** | 5.17 | 7.27 | 1.50 | 0.32 | 0.50 | 0.16 |
| **SNV total[d]** | **108.83** | **61.13** | **83.33** | **61.79** | **81.25** | **85.78** |
| **% SNV[e]** | **80.22** | **81.63** | **72.15** | **61.81** | **92.86** | **81.87** |
| **A/T-1** | 10.83 | 3.77 | 15.00 | 22.92 | 4.00 | 18.02 |
| **G/G-1** | 6.50 | 2.70 | 11.17 | 11.07 | 1.25 | 0.30 |
| **>1 bp del** | 4.00 | 1.19 | 0.50 | 0.09 | 0 | 0 |
| **Deletion total[f]** | **21.33** | **7.66** | **26.67** | **34.07** | **5.25** | **18.31** |
| **% Deletion[g]** | **15.73** | **10.22** | **23.09** | **34.09** | **6.00** | **17.48** |
| **A/T+1** | 3.33 | 4.77 | 4.50 | 3.83 | 1.00 | 0.69 |
| **G/G+1** | 0 | 0 | 0 | 0 | 0 | 0 |
| **>1 bp insert** | 1.00 | 0.75 | 0.83 | 0.25 | 0 | 0 |
| **Insertion total[h]** | **4.33** | **5.53** | **5.33** | **4.07** | **1.00** | **0.69** |
| **% Insertion[i]** | **3.19** | **7.38** | **4.62** | **4.08** | **1.14** | **0.66** |
| **MNV** | 0.83 | 0.50 | 0.17 | 0.02 | 0 | 0 |
| **Replacement** | 0.33 | 0.07 | 0 | 0 | 0 | 0 |
| **Complex total[j]** | **1.17** | **0.57** | **0.17** | **0.02** | **0** | **0** |
| **% Complex[k]** | **0.86** | **0.76** | **0.14** | **0.02** | **0** | **0** |
| **Sum total[l]** | **135.67** | **74.88** | **115.50** | **99.95** | **87.50** | **83.80** |
| **Counts/Freq.** | **1.81** | | **1.16** | | **1.04** | |

| Table S10 (cont'd). Mutation spectra | | | | | | | | |
|---|---|---|---|---|---|---|---|---|
| | *rnr1Y285A* (5)[a] | | *rnr1Y285A msh2Δ* (7) | | *rnr1Y285A msh3Δ* (4) | | *rnr1Y285A msh6Δ* (5) | |
| **Variants** | **Counts[b]** | **Freq.[c]** | **Counts[b]** | **Freq.[c]** | **Counts[b]** | **Freq.[c]** | **Counts[b]** | **Freq.[c]** |
| **CG>AT** | 27.40 | 38.48 | 36.29 | 44.68 | 3.25 | 1.40 | 38.60 | 1 67.69 |
| **CG>GC** | 1.20 | 0.32 | 0.14 | 0.04 | 0 | 0 | 0 | 0 |
| **CG>TA** | 34.00 | 13.68 | 36.43 | 25.48 | 20.25 | 3.23 | 33.40 | 25.62 |
| **TA>AT** | 1.40 | 3.38 | 0.43 | 0.24 | 0 | 0 | 1.20 | 0.78 |
| **TA>CG** | 1.00 | 0.27 | 0.86 | 0.26 | 1.25 | 0.18 | 4.00 | 2.90 |
| **TA>GC** | 2.00 | 2.45 | 0.14 | 0.03 | 0 | 0 | 4.60 | 3.25 |
| **SNV total[d]** | **67.00** | **58.57** | **74.29** | **70.72** | **24.75** | **4.82** | **81.80** | **99.24** |
| **% SNV[e]** | **79.95** | **60.41** | **76.36** | **62.3** | **58.58** | **4.71** | **88.72** | **93.08** |
| **A/T-1** | 4.20 | 9.38 | 9.71 | 23.59 | 4.50 | 0.97 | 8.80 | 7.06 |
| **G/G-1** | 10.20 | 27.98 | 9.57 | 16.61 | 13.00 | 96.45 | 0.80 | 0.18 |
| **>1 bp del** | 0.80 | 0.27 | 0 | 0 | 0 | 0 | 0 | 0 |
| **Deletion total[f]** | **15.20** | **37.63** | **19.29** | **40.20** | **17.50** | **97.42** | **9.60** | **7.24** |
| **% Deletion[g]** | **18.14** | **38.81** | **19.82** | **35.43** | **41.42** | **95.29** | **10.41** | **6.79** |
| **A/T+1** | 1.40 | 0.73 | 3.00 | 2.13 | 0 | 0 | 0.80 | 0.15 |
| **G/G+1** | 0 | 0 | 0.14 | 0.05 | 0 | 0 | 0 | 0 |
| **>1 bp insert** | 0.20 | 0.03 | 0.57 | 0.36 | 0 | 0 | 0 | 0 |
| **Insertion total[h]** | **1.60** | **0.75** | **3.71** | **2.54** | **0** | **0** | **0.80** | **0.15** |
| **% Insertion[i]** | **1.91** | **0.78** | **3.82** | **2.24** | **0** | **0** | **0.87** | **0.14** |
| **MNV** | 0 | 0 | 0 | 0 | 0 | 0 | 0 | 0 |
| **Replacement** | 0 | 0 | 0 | 0 | 0 | 0 | 0 | 0 |
| **Complex total[j]** | **0** | **0** | **0** | **0** | **0** | **0** | **0** | **0** |
| **% Complex[k]** | **0** | **0** | **0** | **0** | **0** | **0** | **0** | **0** |
| **Sum total[l]** | **83.80** | **96.95** | **97.29** | **113.46** | **42.25** | **102.24** | **92.20** | **106.62** |
| **Counts/Freq** | **0.86** | | **0.86** | | **0.41** | | **0.87** | |

| Table S10 (cont'd). Mutation spectra | | | | | | | | |
|---|---|---|---|---|---|---|---|---|
| | *rnr1Y285A pGAL-RNR1* (6)[a] | | *rnr1Y285A pGAL-RNR1 msh2Δ* (4) | | *rnr1Y285A pGAL-RNR1 msh3Δ* (4) | | *rnr1Y285A pGAL-RNR1 msh6Δ* (4) | |
| **Variants** | **Counts[b]** | **Freq.[c]** | **Counts[b]** | **Freq.[c]** | **Counts[b]** | **Freq.[c]** | **Counts[b]** | **Freq.[c]** |
| **CG>AT** | 26.33 | 31.68 | 25.75 | 30.55 | 3.25 | 3.61 | 21.50 | 41.39 |
| **CG>GC** | 1.17 | 0.23 | 1.25 | 0.18 | 0 | 0 | 0.50 | 0.11 |
| **CG>TA** | 33.17 | 8.73 | 41.00 | 18.80 | 18.00 | 3.57 | 17.75 | 7.64 |
| **TA>AT** | 1.83 | 2.26 | 1.00 | 0.25 | 0.50 | 0.19 | 2.00 | 2.01 |
| **TA>CG** | 2.00 | 0.31 | 3.50 | 0.93 | 0.75 | 0.12 | 0.75 | 0.12 |
| **TA>GC** | 1.50 | 0.75 | 3.00 | 3.21 | 0 | 0 | 2.00 | 0.66 |
| **SNV total[d]** | **66.00** | **43.96** | **75.50** | **53.91** | **22.50** | **7.49** | **44.50** | **51.93** |
| **% SNV[e]** | **75.29** | **44.59** | **71.89** | **53.22** | **56.96** | **7.45** | **76.72** | **49.20** |
| **A/T-1** | 7.00 | 3.40 | 15.25 | 29.66 | 3.50 | 1.39 | 1.75 | 1.98 |
| **G/G-1** | 12.00 | 50.42 | 9.75 | 9.03 | 13.50 | 91.61 | 10.25 | 50.09 |
| **>1 bp del** | 0.33 | 0.20 | 0.075 | 0.19 | 0 | 0 | 0.50 | 0.75 |
| **Deletion total[f]** | **19.33** | **54.02** | **25.75** | **38.88** | **17.00** | **93.00** | **12.50** | **52.81** |
| **% Deletion[g]** | **22.05** | **54.79** | **24.18** | **38.38** | **43.04** | **92.55** | **21.55** | **50.03** |
| **A/T+1** | 1.833 | 0.45 | 4.25 | 6.00 | 0 | 0 | 1.00 | 0.81 |
| **G/G+1** | 0 | 0 | 0 | 0 | 0 | 0 | 0 | 0 |
| **>1 bp insert** | 0.17 | 0.05 | 1.00 | 2.50 | 0 | 0 | 0 | 0 |
| **Insertion total[h]** | **2.00** | **0.50** | **5.25** | **8.50** | **0** | **0** | **1.00** | **0.81** |
| **% Insertion[i]** | **2.28** | **0.511** | **4.93** | **8.40** | **0** | **0** | **1.72** | **0.77** |
| **MNV** | 0.17 | 0.05 | 0 | 0 | 0 | 0 | 0 | 0 |
| **Replacement** | 0.17 | 0.05 | 0 | 0 | 0 | 0 | 0 | 0 |
| **Complex total[j]** | **0.33** | **0.10** | **0** | **0** | **0** | **0** | **0** | **0** |
| **% Complex[k]** | **0.38** | **0.10** | **0** | **0** | **0** | **0** | **0** | **0** |
| **Sum total[l]** | **87.67** | **98.58** | **106.50** | **101.28** | **39.50** | **100.48** | **58.00** | **105.55** |
| | **0.89** | | **1.05** | | **0.39** | | **0.55** | |
